# Enhanced tRNA array method version 2 for simultaneous in vitro synthesis of 21 tRNAs

**DOI:** 10.64898/2026.01.06.697847

**Authors:** Ryota Miyachi, Anna Irie, Norikazu Ichihashi

## Abstract

Transfer RNAs (tRNAs) play an essential role in translation, and their simultaneous *in vitro* synthesis remains a key challenge in bottom-up synthetic biology. We previously developed the tRNA array method that enables the simultaneous *in vitro* synthesis of 21 tRNAs from a single DNA template; however, the translational activity was substantially lower than that achieved using individually prepared 21 tRNAs for some proteins. Here, we identify the tRNA groups (PIEN group) that limit translation in the tRNA array method and improve the translational activities through sequence modification and incorporation of a leader sequence into the array construct. The resulting tRNA array method version 2 produces a tRNA set that allows translation at levels similar to those achieved with individually prepared tRNAs for multiple reporter proteins under both translation-coupled and uncoupled conditions. This tRNA synthesis scheme provides an improved platform for constructing self-reproducible gene expression systems.

## Introduction

Living systems possess the ability to reproduce the molecular components that constitute themselves. In vitro reconstitution of a minimal molecular system that exhibits this property is one of the central challenges in bottom-up synthetic biology(1–9). In particular, the construction of a self-reproductive central dogma system is an essential requirement for sustaining life-like systems. Although many components, including translation proteins(10–18), ribosomal components(19–21), transfer RNAs (tRNA)(22–24), and DNA replication machinery(25–31), have been successfully synthesized in an *E. coli*-based reconstituted gene expression system (PURE system)(32) for self-regeneration, the efficiency of these syntheses remains a major challenge. Among these components, tRNA poses a large obstacle(33), as its synthesis requires multiple processing steps that have not been fully reconstituted *in vitro*.

In recent years, the use of *in vitro*-synthesized tRNAs in reconstituted gene expression systems has advanced substantially, enabling a wide range of applications, including genetic code expansion or engineering(33–36). In addition, two research groups, including ours, have demonstrated that functional tRNAs can be expressed within the PURE system and coupled directly with translation reactions(22–24). Li *et al*. achieved sustained translation under a microfluidic chemostat system by expressing 21 tRNAs using nicked DNA templates(24). We also succeeded in the simultaneous expression of 21 tRNAs from a single polycistronic DNA template in the PURE system by combining the self-cleaving HDV ribozyme (HDVR)(37) and RNase P digestion, which we named the tRNA array method(23). While this method provided a useful means for both genetic codon engineering and constructing a self-reproducible translation system, there remains a shortcoming: the translation level with tRNA prepared by this method was significantly lower than that with individually prepared tRNAs for some proteins.

In this study, we aimed to improve the tRNA array method to achieve translation efficiencies comparable to those obtained with individually prepared tRNAs for multiple proteins. To this end, we first identified the tRNA groups that limited translation using the original tRNA array method. We then improved the activities and expression levels of the insufficient tRNAs by introducing mutations to stabilize the cloverleaf structure and introducing leader sequences to enhance processing efficiency. By integrating these approaches, we succeeded in enhancing the expression levels of at least three proteins to levels comparable to those obtained with individually prepared tRNA mixtures under both translation-uncoupled and -coupled conditions.

## Materials and Methods

### DNA preparation

DNA templates for preparation of tRNA variants were generated by PCR using forward oligonucleotides containing a T7 promoter and the corresponding tRNA sequences, together with 2’-O-methyl–modified reverse primers (Table S1). The combinations of tRNA variant sequences and primers used in this study are listed in Table S2. DNA fragment encoding L_PIEN_v2 was obtained using Twist Bioscience’s artificial gene synthesis service. Replacement of the original PIEN region in the pTwist_21tRNAs plasmid with the improved PIEN sequences was performed by amplifying the corresponding fragments using specific primers (primers 20 and 21 for the insert fragment and primers 22 and 23 for the vector fragment) and assembling them using In-Fusion cloning kit (Takara). The resulting DNA sequences are listed in Table S3. A linear DNA template encoding all 21 tRNAs (a single 21 tRNA template) was amplified by PCR using primers 24 and 21 for use in expression-coupled translation experiments.

The luciferase template DNA was prepared by PCR amplification using primers 25 and 26 (Table S1) and a plasmid (pUC-T7p-Fluc_21tRNA) constructed in our previous study(22), followed by column purification. DNA fragments encoding sfGFP and GUS were obtained using an artificial gene synthesis service (Twist Bioscience), as described previously(23), and used for DNA templates used for translation experiments in the PURE system after PCR amplification using the same primer pair (primers 25 and 26) and column purification. The sequences of all reporter gene templates are listed in Table S3. All DNA concentrations were quantified based on the A_260_.

### Individual tRNA and M1 RNA preparation

The sequences of the 21 tRNAs used in this study were based on our previous work(22,23) and are listed in Table S4. Individually-prepared IVS tRNAs were synthesized essentially as described previously. Briefly, among the 21 tRNAs, 15 species initiating with guanosine at the 5′ end were individually prepared by *in vitro* transcription whereas the remaining six non-G-start tRNAs were chemically synthesized (Invitrogen), as in our earlier studies. The DNA template for M1 RNA was prepared by PCR using a plasmid encoding M1 RNA as described previously and primers 27 and 28 (Table S3).

*In vitro* transcription was performed in a mixture containing 40 mM Tris-HCl pH 8.0, 10 mM magnesium chloride, 2 mM spermidine, 5 mM DTT, 1 U/μl T7 RNA polymerase (Takara), 2 mM each NTPs(ATP, GTP, CTP, and UTP), 3 mM GMP, 5 μg/μl template DNA, 2 U/μl inorganic pyrophosphatase (New England Biolab), 0.8 U/μl RNasin (Promega). The mixture was incubated at 37°C for 12 h, and the RNA product was individually purified using the PureLink RNA Mini Kit (Invitrogen). Both tRNAs and M1 RNA were dissolved in water and stored at -80 °C until use. RNA concentrations were determined based on the A260.

### Simultaneous 21 tRNAs preparation using the tRNA array method

Simultaneous synthesis of 21 tRNAs was performed using the tRNA array method with a single linear DNA template encoding all 21 tRNAs (25 nM). *In vitro* transcription was carried out with the single 21 tRNAs template (25 nM) in the presence of Tris-HCl pH 8.0 (40 mM), magnesium chloride (10 mM), spermidine (2 mM), T7 RNA polymerase (1.0 U/μL, Takara) and T4 PNK (0.1 U/μL, New England Biolab), NTPs (2 mM each: ATP, GTP, CTP, and UTP), GMP (3 mM), 0.2 U/μl inorganic pyrophosphatase (New England Biolab), and 0.8 U/μl RNasin (Promega) at 37°C for 12 h. The resulting array-derived pre-tRNA transcripts were purified using the PureLink RNA Mini Kit (Invitrogen). The purified premature tRNA transcripts were subsequently processed by RNase P digestion in buffer R (see RNase P digestion in buffer R section) with RNase P (4 μM M1 RNA and 6 μM C5 protein) at 37 °C for 12 h for digestion. Following processing, the resulting tRNA products were purified again using the PureLink RNA Mini Kit and used for subsequent translation assays. C5 protein was purified as described previously(22,23)

### tfPURE system preparation

All components of the laboratory-prepared PURE system were expressed and purified essentially as described previously, using affinity chromatography with a histidine tag, followed by gel-filtration chromatography (38). EF-Tu was further purified by two rounds of affinity chromatography in a stringent buffer to remove residual tRNA, as previously reported(22). Ribosomes were further purified by ultrafiltration to remove residual tRNAs, following the same procedure used in our previous work. The detailed composition of the tfPURE system used in this study is summarized in Table S5. The composition A is the same composition we used in our previous study(22). The compositions B and C are the compositions optimized for four and 21 tRNA expressions, respectively, in our previous study (22,23).

### tRNA design

Secondary structure prediction of each tRNA variant was performed using the RNAfold Web Server from the ViennaRNA package(39). For each sequence, both the minimum free energy (MFE) structure and the centroid structure were obtained, and formation of the canonical cloverleaf structure was evaluated. A cloverleaf structure was defined by the presence of a well-formed acceptor stem, D arm, anticodon arm, and T arm in the predicted secondary structure. In addition, ensemble diversity and the frequency of the MFE structure within the structural ensemble were calculated to assess structural stability and heterogeneity. These parameters were used to compare predicted secondary structures and to select tRNA variants for experimental validation.

### RNase P digestion in buffer R

RNase P digestion of pre-tRNA transcripts was performed in buffer R, consisting of 50 mM Tris-HCl (pH 7.6), 60 mM NH_4_Cl, 10 mM Mg(OAc)_2_, and 5 mM spermidine. Pre-tRNA substrates (250 nM) were incubated with M1 RNA (500 nM) and C5 protein (750 nM) in buffer R at 37 °C for 12 h.

### Urea-PAGE

Denaturing polyacrylamide gel electrophoresis was performed using 8% polyacrylamide gels (acrylamide: bisacrylamide = 19:1) containing 8 M urea, 0.1% ammonium persulfate, and 0.1% N, N, N’, N’-tetramethylethylene-diamine in Tris-borate-EDTA buffer. RNA samples were mixed with stripping buffer composed of 50 mM EDTA, 90% formamide, and 0.025% bromophenol blue prior to electrophoresis. Following electrophoresis, RNA was visualized by staining with SYBR Green II (Takara). An RNA size marker (Dyna Marker RNA Low II Easy Load, Bio Dynamics Laboratory Inc.) was loaded alongside samples in all urea-PAGE analyses.

### RT-qPCR for tRNA quantification

For quantification of individual tRNA species, 21-tRNA mixtures prepared by the tRNA array method were purified by size selection using the RNA Clean & Concentrator system (Zymo Research) to enrich the tRNA fraction. The purified 21-tRNA mixtures were diluted to 500 pM as the total 21 tRNAs and analyzed by RT-qPCR using the One Step TB Green PrimeScript PLUS RT-PCR Kit (Takara) with tRNA-specific primer sets (Supplementary Table S1). The following primer pairs were used: Gly, primers 29 and 30; Pro_v1, primers 31 and 32; Pro_v2, primers 33 and 32; Ile_v1, primers 34 and 35; Ile_v2, primers 36 and 35; Glu, primers 37 and 38; Asn_v1, primers 39 and 40; and Asn_v2, primers 41 and 40. Reactions were performed on a QuantStudio 3 Real-Time PCR System (Thermo Fisher Scientific) under the following conditions: reverse transcription at 42 °C for 30 min; initial denaturation at 95 °C for 10 sec; followed by 50 cycles of 95 °C for 5 sec and 60 °C for 30 sec, with fluorescence measured at the end of each cycle.

For quantification, standard curves were generated using serial dilutions of individually prepared tRNAs with known concentrations, and the concentrations of the corresponding tRNAs in the 21-tRNA mixtures were determined from the measured Ct values. Primer specificity was evaluated by comparing amplification of the target tRNA with that of the corresponding non-target 20-tRNAs mixture.

### Translation with purified 21 tRNAs

Translation reactions were carried out in the laboratory-prepared tRNA-free PURE system (tfPURE, composition A, Table S6) supplemented with a prepared 21-tRNA mixture, a reporter gene encoding DNA template (1 nM). The reaction mixture was incubated at 30 °C for 16 h. The total concentration of the individually prepared 21 tRNAs was 240 ng/μL (40 ng/μL each for PIEN and 4.7 ng/μL each for the others). When 21 tRNAs prepared by the tRNA array method were used, the total tRNA concentration was also set to 240 ng/μL. For assays involving tRNA variants, PURE systems lacking the respective tRNA were prepared.

After incubation, reporter protein production was quantified using three independent assays. For luciferase, a 1 μL aliquot of the translation reaction was mixed with 30 μL of Luciferase Assay Reagent (Promega), and luminescence was measured using a GloMax luminometer (Promega). For sfGFP, fluorescence was monitored for 24 h using an Mx3005P (Agilent Technologies). For GUS, a 2 μL aliquot of the translation reaction was added to 8 μL of GUS reaction buffer (final concentrations: 50 mM HEPES-KOH, pH 7.4; 0.01% Triton X-100; 5 μM TokyoGREEN-βGlcU(Na); Sekisui Medical, Japan)(40). Fluorescence was recorded every 1 min for 60 min using an Mx3005P with FAM detection settings. GUS activity was calculated from the slope of the linear portion of the fluorescence time course.

### Translation coupled with tRNA expression

tRNA-expression coupled translation reactions were performed in the tfPURE system (composition B or C, Table S6) by addition of reporter gene encoding DNA templates (1 nM) and tRNA template DNA. The concentrations of tRNA template DNA, T7 RNA polymerase (Takara), T4 PNK (New England Biolab), and RNase P were adjusted depending on the experimental design and are specified in the corresponding figure legends. The reaction mixture was incubated at 30°C for 24 h. After incubation, translation outputs were quantified using reporter-specific assays for luciferase, sfGFP, or GUS, as described above.

## Results

### Identification of limiting tRNAs

In our previous work, we constructed a tRNA array method that encodes all 21 tRNA species required for translation, which are transcribed from five polycistronic gene groups, in which each tRNA gene in the groups is directly linked in an end-to-end manner (Fig. 1A). After transcription, the five transcripts are processed through self-cleavage of HDVR at the 3’-end and digestion with RNase P to generate mature-length tRNA products. The translation level of luciferase using the 21 tRNAs synthesized by this method was at a similar level to that using individually prepared *in vitro*-synthesized (IVS) tRNA mixtures of all 21 species, but was significantly lower when using sfGFP and beta-glucuronidase (GUS) as reporter genes(23).

**Figure 1.**
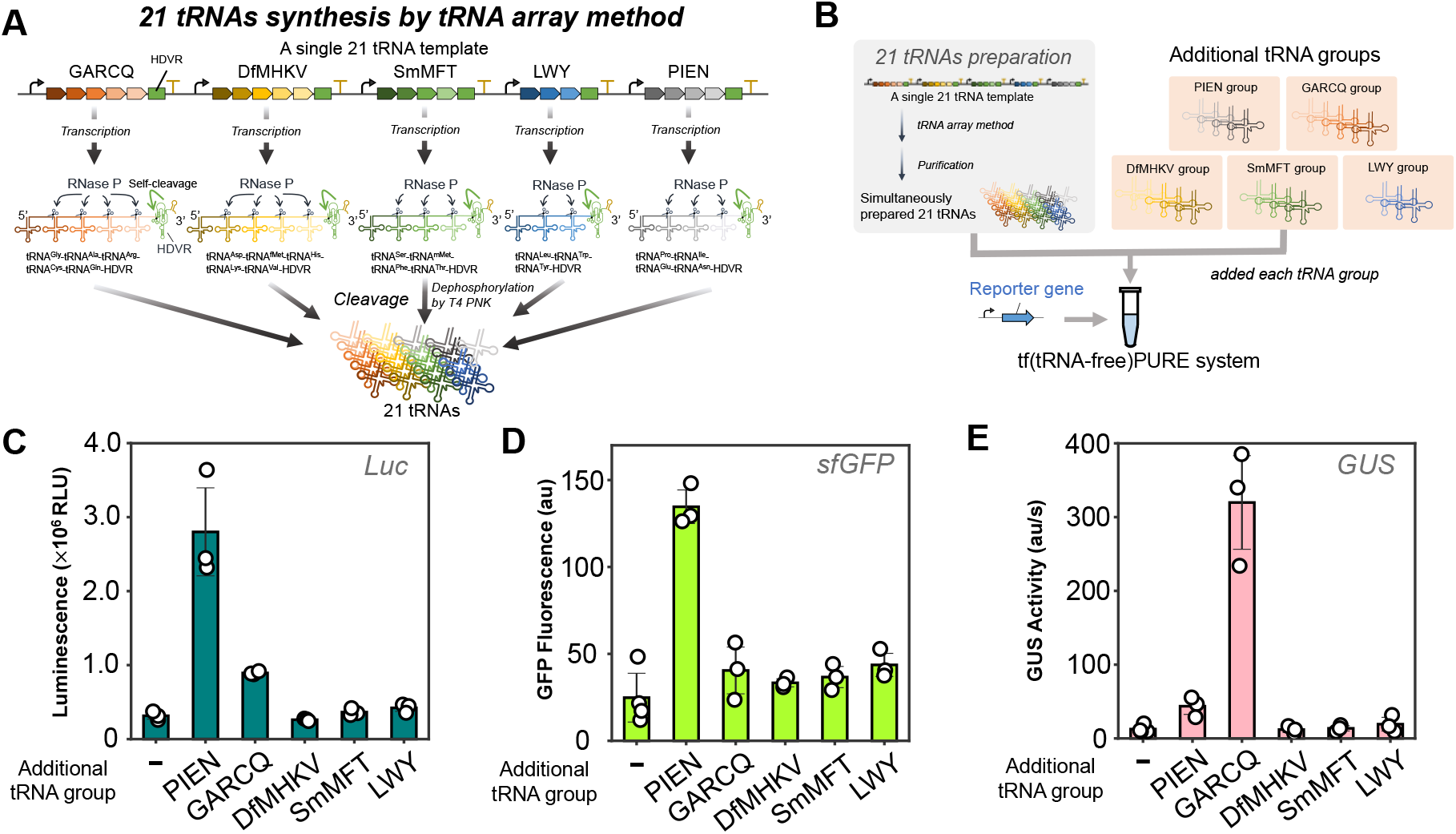
Identification of limiting tRNA groups. (A) Schematic of 21 tRNAs synthesis by the tRNA array method. The 21 tRNA genes are divided into five groups and encoded in a single DNA template. After transcription, five transcripts are processed into mature-sized tRNAs through HDVR self-cleavage and RNase P digestion. (B) Schematic of the supplementation assay. The purified 21 tRNAs (240 ng/μL) synthesized by the tRNA array method and an individually-synthesized additional tRNA mixture (PIEN: 40 ng/μL each; others: 4.7 ng/μL each), corresponding to one of the five groups, were used for translation of three reporter proteins (firefly luciferase, sfGFP, and GUS) in a tRNA-free PURE system (tfPURE; composition A, see Tables S5 and S6) containing reporter-encoding DNA template (1 nM) and T7 RNA polymerase (0.42 U/μL) at 30 °C for 16 h. (C) Luciferase activities. (D) sfGFP fluorescence. (E) GUS activities. GUS activity was quantified by an enzymatic assay after the reaction, and the slope of fluorescence increase over time was used as a measure of activity. Error bars indicate standard deviations (N = 3).

To examine whether the reduced translation level was caused by insufficiency of some tRNAs, we performed a supplementation assay (Fig. 1B), in which we added five individually-prepared tRNA mixtures to the 21 tRNA mixtures prepared by the tRNA array method and compared the translation efficiency of three reporter proteins (luciferase, sfGFP, and GUS) in a tRNA-free PURE system (tfPURE). The compositions of the five added tRNA mixtures were based on the five array groups (GARCQ, DfMHKV, SmMFT, LWY, and PIEN; see Fig. 1A). The total concentration of the 21 tRNAs was set to 240 ng/μL, under which, translation output increased linearly with tRNA concentration (Fig. S1). The concentration of each supplemented tRNA was 40 ng/μL for PIEN tRNAs (tRNA^Pro, Ile, Glu, Asn^) and 4.7 ng/μL for the others, which was the same relative composition as in the reconstituted 21 tRNAs system reported previously(22,33), but uniformly scaled down to 40% of the original concentrations.

Under these conditions, supplementation with the PIEN tRNA mixture increased translation by approximately 3- to 11-fold for all three reporter proteins: firefly luciferase, sfGFP, and GUS. In contrast, supplementation with the GARCQ tRNAs enhanced translation of luciferase (approximately three-fold) and GUS (approximately 20-fold), but did not increase sfGFP translation, indicating that the optimal tRNA composition varies depending on the protein to some extent. Because the addition of PIEN tRNAs consistently enhanced translation across all three proteins, we focused subsequent optimization efforts on this group.

### Sequence modification of tRNA^Ile^, tRNA^Asn^, and tRNA^Pro^

We attempted to improve the translational activity of tRNAs in the PIEN group. Previous studies have shown that stabilization of the canonical cloverleaf structure can enhance tRNA functionality(41,42). Secondary structure prediction revealed that the tRNA^Ile^ and tRNA^Asn^ used here, which are the same as *E. coli* native sequences, do not form standard cloverleaf structures as their minimum free energy (MFE) structures (Fig. S2). In addition, while the MFE structure of tRNA^Pro^ is predicted to form a cloverleaf, the anticodon stem region exhibits reduced structural stability (Fig. S2). These non-standard structures may impair their translational activities. We therefore introduced mutations to tRNA^Ile^, tRNA^Asn^, and tRNA^Pro^ to adopt stable cloverleaf structures.

To make stable cloverleaf structures, we first introduced point mutations primarily into loop regions while avoiding identity elements(43) and generated various tRNA variants. We then predicted the secondary structures of the resulting variants using the ViennaRNA package(39) and evaluated the frequency at which the MFE structure formed the canonical cloverleaf conformation and the ensemble diversity, which quantifies the structural heterogeneity of the RNA ensemble. Variants exhibiting a high MFE frequency and low ensemble diversity were selected for the following translation assay. In the translation assay, tfPURE was supplemented with either a tRNA variant or the original tRNA, together with a mixture of the remaining 20 tRNAs and DNA encoding luciferase. The transcription and translation reactions were carried out by incubating the mixture at 30 °C for 16 h, and the luciferase activity was measured.

For tRNA^Ile^ and tRNA^Asn^, we selected five point mutants that exhibit increased frequencies of the cloverleaf structure for translation assay (Figs. 2A and 2C). The percentage of MFE structures that adopt a cloverleaf conformation are shown below in blue. For tRNA^Ile^, three out of five tested variants exhibited similar translation activity to that of the original tRNA (Fig. 2B). For tRNA^Asn^, three of the five variants showed similar activities to that of the original tRNA (Fig. 2D). In contrast, some tRNA variants (C32G and U33A in tRNA^Ile^, and C32G and A38G in tRNA^Asn^), which contain mutations in the anticodon loop, significantly reduced translation activity, indicating that these regions are essential for translation activity, although they have not been reported as identity elements.

**Figure 2.**
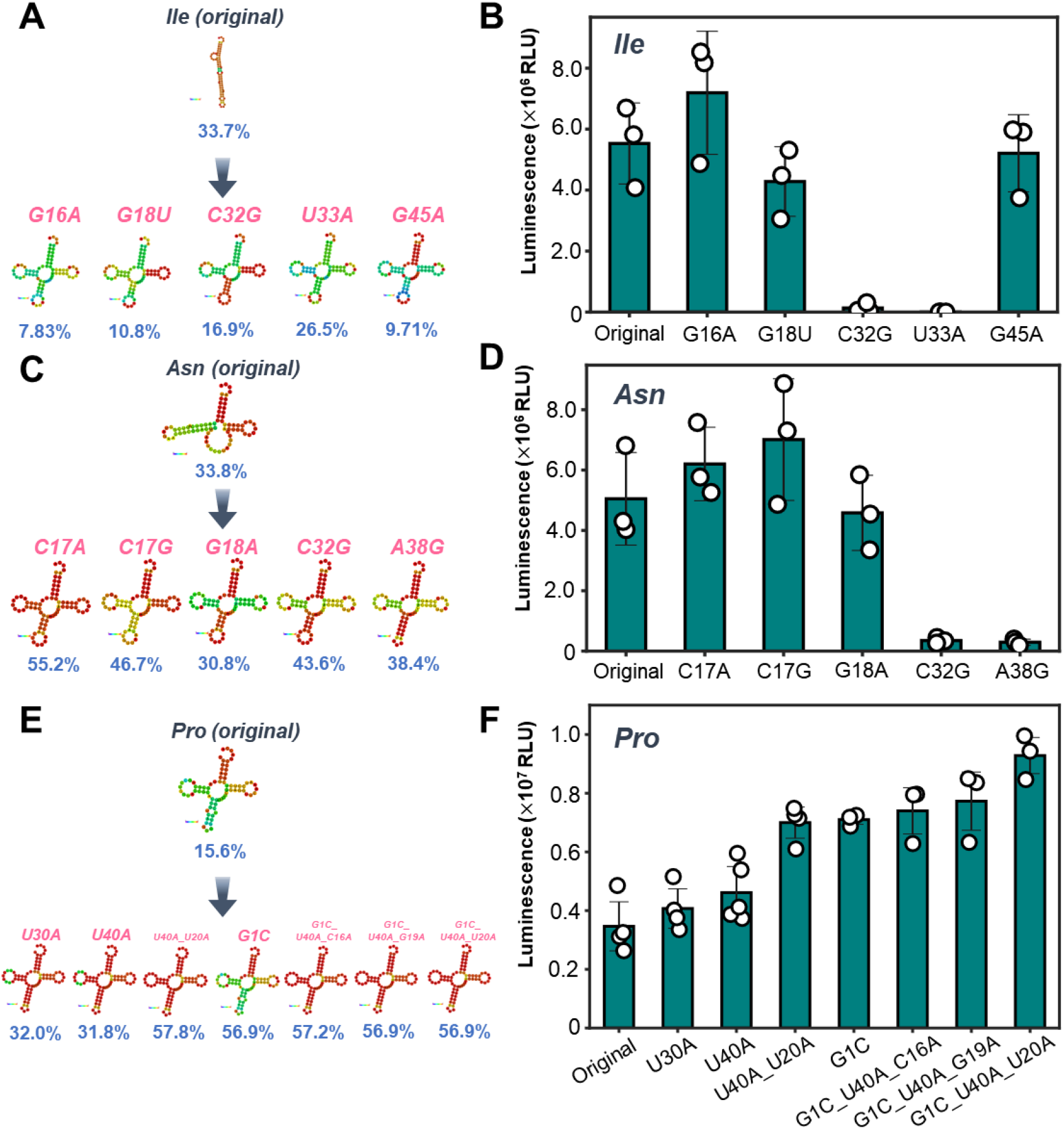
Sequence modification of tRNAs to stabilize cloverleaf structures. (A, C, E) Minimum free energy (MFE) secondary structures of tRNA^Ile^(A), tRNA^Asn^(C), and tRNA^Pro^ (E) variants predicted using ViennaRNA(39). The percentages indicate the frequency with which the corresponding minimum free energy (MFE) structure was predicted to adopt the canonical cloverleaf conformation among the ensemble. The number of mutational site follows the universal conventional tRNA positions (i.e., anticodons are 34–36 for all tRNAs). (B, D, F) Translation activity assays of tRNA^Ile^ (B), tRNA^Asn^ (D), and tRNA^Pro^ (F) variants. Translation reactions were performed in the tfPURE system (composition A) containing a 20-tRNAs mixture (500 ng/μL) lacking the target tRNA, one of the target tRNA variants (tRNA^Ile^ or tRNA^Asn^ at 13 ng/μL, tRNA^Pro^ at 25 ng/μL), a DNA template encoding luciferase (1 nM), and T7 RNA polymerase (0.42 U/μL). Reactions were incubated at 30 °C for 16 h and luciferase activity was measured. Error bars indicate standard deviations (N = 3).

For tRNA^Pro^, we selected seven sequences that exhibit higher frequencies of cloverleaf conformation (Fig. 2E). In these variants, we additionally introduced a mutation (G1C) at the 5’-terminus. The original nucleotide at this position in wild-type tRNA^Pro^ is C, but was mutated to G in the previous array method to allow transcription to start immediately downstream of the T7 promoter(23). To examine the possibility that this mutation could impair translation activity, we selected four tRNA^Pro^ variants that reverted this position to C (G1C). Among these variants, the G1C_U40A_U20A variant exhibited the highest translation activity, reaching approximately 2.5-fold higher activity than the original tRNA (Fig. 2F).

### Evaluation of PIEN array arrangement

We next examined the arrangement of the PIEN group on the DNA template. In the tRNA array method, transcription generates premature transcripts that are first processed by HDVR self-cleavage at the 3’-end. In our previous study, we observed that the PIEN group exhibited relatively low efficiency at this self-cleavage step(23). Because this processing step may depend on the order of tRNA genes within the array, we constructed four PIEN group arrays with different tRNA orders and synthesized the corresponding premature transcripts in the absence of RNase P to directly compare HDVR self-cleavage efficiencies (Fig. 3A). The self-cleavage efficiency varied among these four constructs, and the original PIEN arrangement showed the highest self-cleavage efficiency (Fig. 3B, blue arrowheads). In addition, the NIPE arrangement exhibited a larger proportion of uncleaved products of larger size. This is likely due to predominant transcription termination at the second stem loop of the tandem T7 terminator(44) for unknown reasons, resulting in longer RNA species.

**Figure 3.**
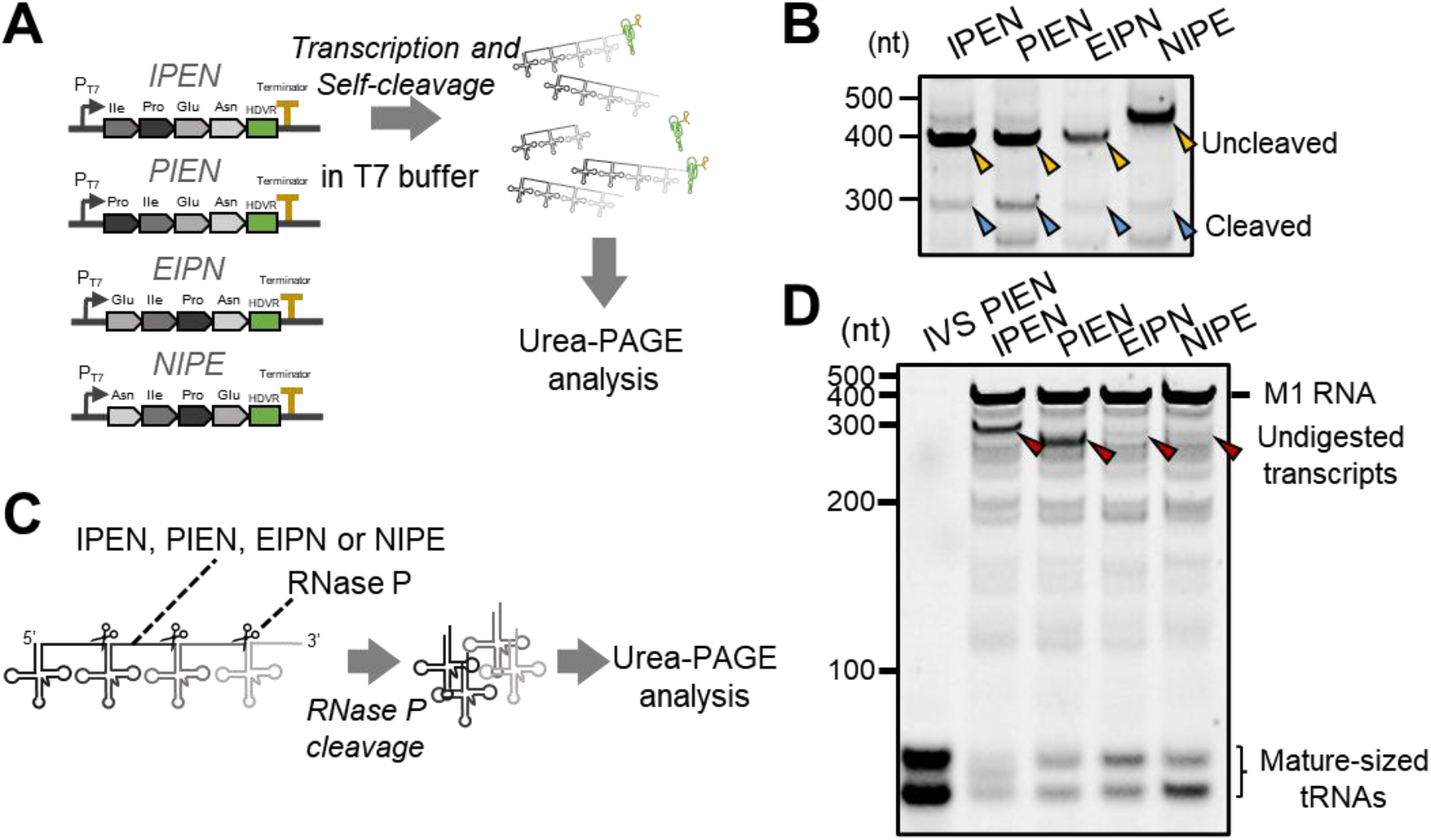
Evaluation of PIEN group arrangement. (A) Schematic of the HDVR self-cleaving assay of PIEN group constructs. Transcription was performed using templates with different tRNA orders in the absence of RNase P, and the resulting transcripts were analyzed by urea-PAGE. (B) Urea-PAGE analysis of transcripts from PIEN group constructs, followed by SYBR Green II staining. Predicted bands corresponding to transcripts containing HDVR and those lacking HDVR are indicated by blue and yellow arrowheads, respectively. (C) Schematic of the experimental workflow for the RNase P digestion assay. Prepared pre-tRNAs (250 nM) lacking HDVR were used as substrates and incubated with RNase P (500 nM M1 RNA and 750 nM C5 protein) in buffer R (see Methods). (D) Urea-PAGE analysis of the digested products, followed by SYBR green II staining. Individually prepared PIEN tRNA mixtures were included as size controls. Undigested pre-tRNA species are indicated by red arrows.

Next, we examined whether the gene arrangement affected the yield of RNase P processing. To enable direct comparison after self-cleavage, pre-tRNAs lacking the HDV ribozyme sequence were prepared by *in vitro* transcription. These pre-tRNAs were incubated with RNase P in buffer R (see Methods) at 37 °C for 12 h, and the cleavage products were analyzed by urea-PAGE (Fig. 3C). The EIPN and NIPE arrangements exhibited the largest products, while the PIEN arrangement exhibited slightly smaller products (Fig. 3D, the bands indicated as “mature-sized tRNAs”). It should be noted that in our previous study (23), we reported translation with these constructs but did not analyze each HDVR and RNase P cleavages. The present analysis was therefore performed to clarify these processing steps.

### PIEN array version 2

Based on these results, we decided to use the original PIEN tRNA order because it exhibited the highest HDVR self-cleavage (Fig. 3B) and modest RNase P digestion (Fig. 3D). We then proceeded to replace the original tRNA sequences in the PIEN array with the cloverleaf-stabilized variants that showed higher translation activities (tRNA^Pro^_G1C_U40A_U20A, tRNA^Ile^_G16A, and tRNA^Asn^_C17G, respectively). Because the T7 RNA polymerase we used starts RNA synthesis from G nucleotides, to adopt the tRNA^Pro^, which has a non-G at the 5’-terminus, we introduced a 27-nucleotide leader sequence containing 5’-G upstream of the tRNA^Pro^_G1C_U40A_U20A gene (Fig. 4A). The first eight nucleotides of the leader sequence were designed for efficient transcription initiation(45) and the last six nucleotides were designed to enhance RNase P cleavage efficiency(46). The intervening sequence was adopted from a previously reported leader sequence(33). The resulting PIEN construct was designated as L_PIEN_v2.

**Figure 4.**
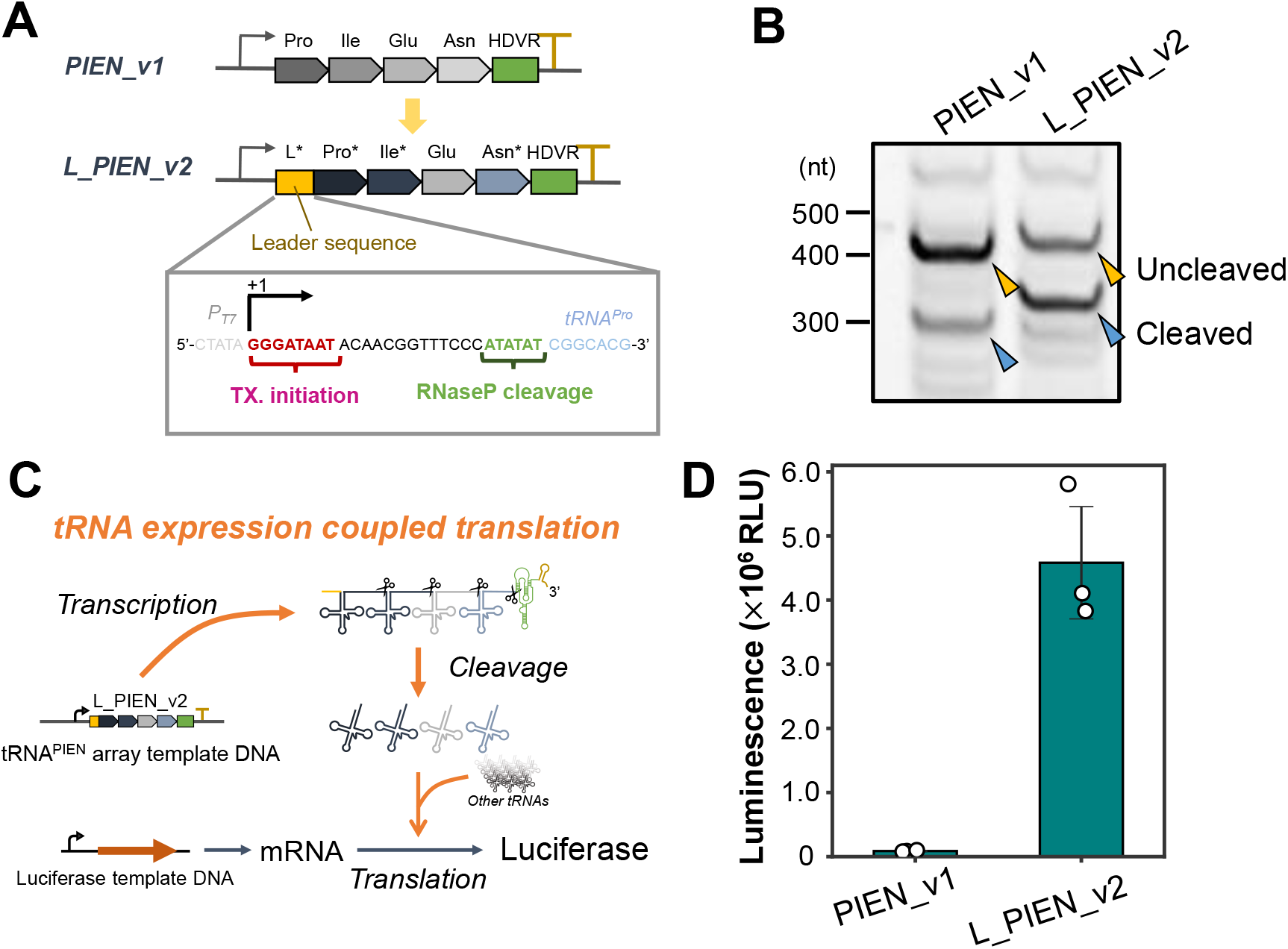
Cleavage of and translation with PIEN array version 2. (A) Design of the improved PIEN construct (L_PIEN_v2), in which tRNA^Pro^, tRNA^Ile^, and tRNA^Asn^ were replaced with the higher-activity variants (tRNA^Pro^_G1C_U40A_C20A, tRNA^Ile^_G16A, and tRNA^Asn^_C17G, respectively) and a leader sequence was inserted upstream of tRNA^Pro^. (B) Urea-PAGE analysis of transcripts generated from the previous construct (PIEN_v1) and L_PIEN_v2 templates, followed by SYBR green II staining. Predicted bands corresponding to the uncleaved transcript containing HDVR and the transcript after self-cleavage of HDVR are indicated by yellow and blue arrowheads, respectively. (C) Schematic of translation assay using PIEN tRNAs. Reactions were performed in tfPURE (composition B) supplemented with the tRNA^PIEN^ array template DNA (25 nM), luciferase template DNA (1 nM), RNase P (1 μM M1 RNA and 1.5 μM C5 protein), T4 polynucleotide kinase (0.094 U/μL), T7 RNA polymerase (1.7 U/μL), and 17 IVS tRNAs that lack PIEN tRNAs. After incubation at 30 °C for 24 h, luciferase activity was measured. (D) Measured luciferase activities. Error bars indicate standard deviations (N = 3).

We first compared the efficiency of HDVR self-cleavage of L_PIEN_v2 with that of the previous construct (PIEN_v1) by performing *in vitro* transcription followed by urea-PAGE analysis (Fig. 4B). Quantification of the ratio of cleaved (blue arrowheads) to uncleaved (yellow arrowheads) products revealed that the original PIEN construct yielded approximately 25% cleaved products, whereas L_PIEN_v2 demonstrated an efficiency of approximately 53%, corresponding to a 2.1-fold improvement. When only the tRNA sequences were replaced without the leader sequence (PIEN_v2), the self-cleavage efficiency was intermediate between that of PIEN_v1 and L_PIEN_v2 (Fig. S3).

We next evaluated the translation activity of the expressed tRNA set by a translation-coupling reaction (Fig. 4C). In this assay, the template DNA encoding the PIEN arrays was used at a concentration of 25 nM, corresponding to half the DNA concentration used in our previous study(23), to evaluate translation activity under an unsaturated condition. Under this condition, the original PIEN construct exhibited a luciferase activity of 8.8×10^4^, whereas L_PIEN_v2 exhibited a value of 4.6×10^6^, representing an approximately 52-fold increase in translation activity. This large improvement may reflect the combined effects of enhanced HDVR processing, improved tRNA function, and the non-proportional increase in translation with increasing PIEN tRNA concentration in this range (Fig. S4).

### Construction and evaluation of tRNA array version 2

L_PIEN_v2 was incorporated into the polycistronic DNA construct encoding all 21 tRNAs used in our previous study as the tRNA array version 2. Using this DNA, we synthesized 21 tRNAs under a translation-uncoupled condition, purified them, and evaluated the translation activity for three reporter genes (luciferase, sfGFP, and GUS) at a total tRNA concentration of 240 ng/μL (Fig. 5A). For luciferase translation, the original tRNA array (v1) yielded a luminescence signal of approximately 4.1×10^5^, whereas tRNA array version 2 (v2) produced a signal of approximately 4.3×10^6^, corresponding to an approximately 11-fold increase (Fig. 5B). Similarly, sfGFP translation was enhanced by approximately 4.1-fold (Fig. 5C) and GUS translation increased by approximately 5.1-fold with tRNA array version 2 (Fig. 5D). Furthermore, for all three reporter genes, tRNA array version 2 supported translation activities comparable to, or exceeding, those achieved with individually prepared 21 IVS tRNA mixtures (IVS) at the same concentration, the composition of which is based on a previous study(33). RT-qPCR analysis of purified 21-tRNA mixtures showed that the abundances of the redesigned PIEN-group tRNAs were increased in version 2 relative to version 1, whereas tRNA^Gly^, the construct of which was unchanged, exhibits a slight decrease (Fig. S5).

**Figure 5.**
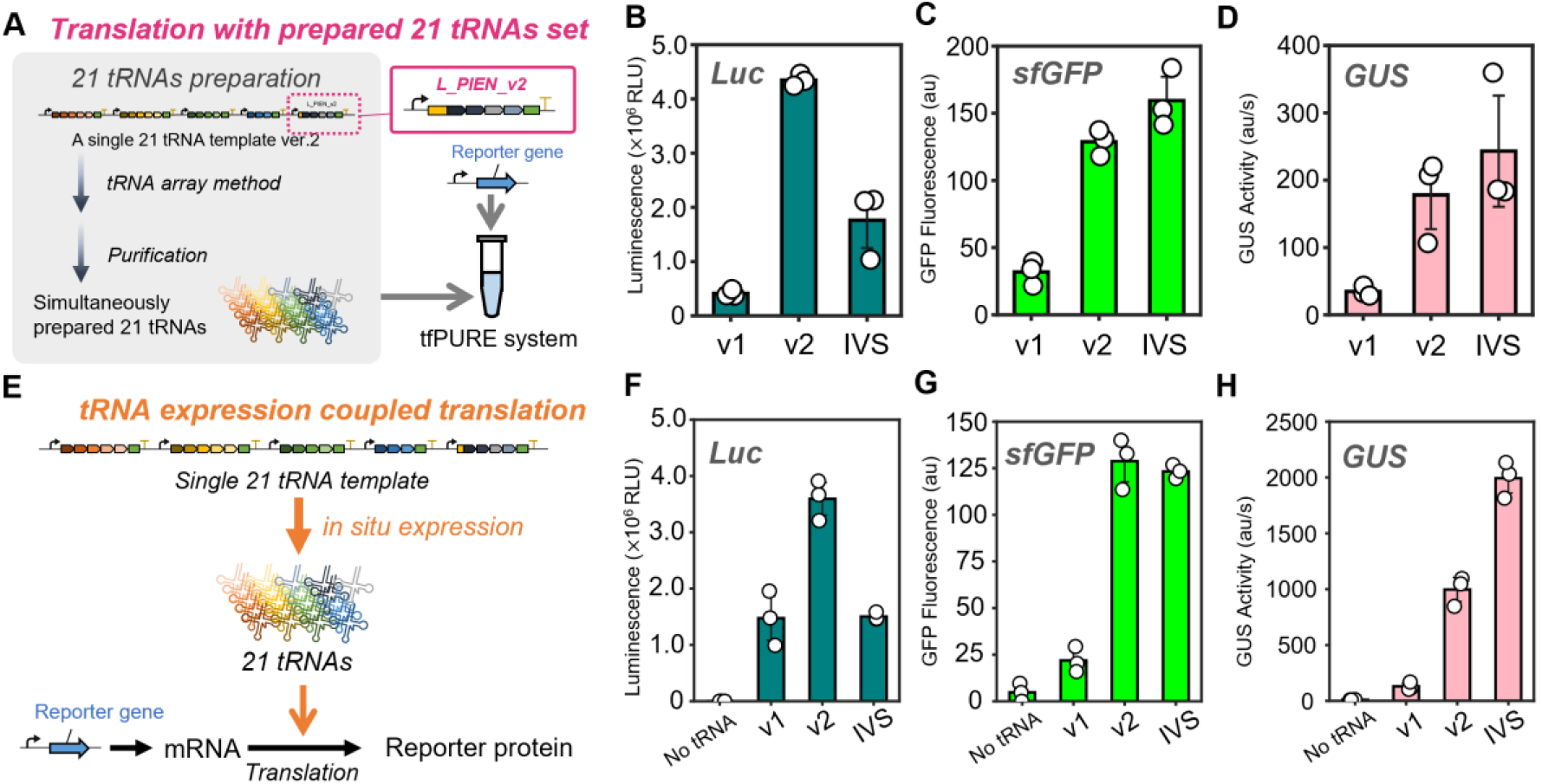
Translation with 21 tRNAs prepared by using tRNA array ver. 2. (A) Schematic of translation assays using 21 tRNAs prepared by the translation-uncoupled method. Twenty-one tRNAs were simultaneously expressed using the tRNA array method from a 21-tRNA array version 2, in which the PIEN group was replaced with version 2 variants. The resulting 21-tRNA mixture (240 ng/μL) was purified and added to tfPURE (composition A) together with a reporter gene (luciferase, sfGFP, or GUS)–encoding DNA template (1 nM) and T7 RNA polymerase (0.42 U/μL), followed by incubation at 30 °C for 16 h before activity measurements. (B) Luciferase activities. (C) sfGFP fluorescence. (D) GUS activity. (E) Schematic of translation-coupled *in situ* synthesis of all 21 tRNAs using tRNA array version 2. Reactions were performed in tfPURE (composition C) supplemented with the 21 tRNA template (50 nM), a reporter gene (luciferase, sfGFP, or GUS)–encoding DNA(1 nM), T7 RNA polymerase (3.4 U/μL), T4 polynucleotide kinase (0.094 U/μL), and RNase P (4 μM M1 RNA and 6 μM C5 protein). When using individually prepared 21 IVS tRNAs (IVS), the concentration was set to 780 ng/μL. After incubation at 30 °C for 24 h, translation activities were measured. (F) Luciferase activities. (G) sfGFP fluorescence. (H) GUS activity. Error bars indicate standard deviations (N = 3).

We compared the performance of tRNA array version 2 in the translation-coupled method. In this experiment, all 21 tRNAs were synthesized in situ within tfPURE and directly utilized for translation (Fig. 5E). First, we measured tRNA synthesis in tfPURE. Quantification of total tRNA production revealed that tRNA array version 2 yielded 780 ng/μl, approximately 1.3-fold higher total tRNA levels than version 1 (Fig. S6). When measuring translated luciferase activity, tRNA array version 2 supported an approximately three-fold higher translation than that with version 1, and an approximately 2.4-fold increase relative to individually prepared 21 IVS tRNA mixtures at the same concentration (Fig. 5F). Similarly, translation of sfGFP was enhanced by approximately six-fold compared with version 1 and reached a level comparable to that obtained with individually prepared 21 IVS tRNAs (Fig. 5G). GUS translation increased by approximately 7.8-fold relative to version 1, reaching approximately 50% of the level achieved with individually prepared 21 IVS tRNAs (Fig. 5H). The absolute GUS activity values differed more markedly between the translation-uncoupled and translation-coupled reactions than those of luciferase or sfGFP, possibly because the two reactions differ substantially in reaction composition and time-dependent tRNA supply. The oligomeric GUS reaction may be more sensitive to these changes.

## Discussion

In this study, we identified the tRNA groups that limited translation activity in the tRNA array method established in our previous work(23), and constructed and developed tRNA array version 2 by improving both tRNA activity and pre-tRNA processing efficiency. Specifically, we identified the PIEN group as a bottleneck for translation of all measured proteins, enhanced the activities of tRNAs (especially tRNA^Pro^) through stabilization of their cloverleaf structures, and redesigned the PIEN array by incorporating a leader sequence. As a result, the translation activity of 21 tRNAs prepared using tRNA array version 2 was improved to levels similar to those achieved with individually prepared tRNAs across multiple reporter proteins, enabling stable expression of all 21 tRNAs in translation-coupled reaction in the PURE system. Our results highlight that quantitative tuning of tRNA composition is critical for efficient translation and represents a key requirement for constructing self-reproducible gene expression systems(47).

Our results demonstrated that translation activity strongly depended on the relative composition of the 21 tRNAs. Translation activity with the previous tRNA array method was increased by supplementation with the PIEN tRNA group for all tested reporter genes (Fig. 1), indicating that reduced translation output from array-derived tRNAs was primarily attributable to insufficient levels of these tRNA species. Consistently, the PIEN tRNA set was used at higher concentrations in a previous study that used an *in vitro*-transcribed tRNA set(33). Considering that the relative expression level of each group was similar, as shown in our previous study(23), this result suggests that efficient translation does not require uniform concentrations of all 21 tRNAs, but requires adjusted concentrations that contain more of the highly-demanded tRNA species, such as PIEN tRNAs.

This study demonstrated that tRNA^Pro^, tRNA^Ile^, and tRNA^Asn^ were predicted to either fail to stably adopt the canonical cloverleaf structure or to exhibit high structural heterogeneity, and some of their variants with more frequent cloverleaf structures exhibited enhanced translation activity. These results suggest that the canonical cloverleaf structure plays a certain role in translation, probably through interactions with translation factors(48,49). Considering that their original sequences are derived from *E. coli* native tRNAs, this result further suggests that native tRNA sequences are not always fully suited to in vitro translation using unmodified tRNAs. However, some mutants (e.g., G18U and G18A in Fig. 2) that stabilized the cloverleaf structures did not enhance translation, indicating that other unknown factors are also important for tRNA function.

The introduction of leader sequences was another key factor contributing to the improved translation performance of tRNA array version 2. Although the leader sequence was designed to allow transcription of tRNA^Pro^ variants that start with 5’-C, an unexpected improvement was observed in HDVR self-cleavage efficiency. This finding suggests that incorporation of leader sequences and modification of tRNA sequences altered the overall secondary structure of the pre-tRNA, potentially generating conformations that are more favorable for HDVR cleavage. In addition, insertion of the leader sequence is expected to mitigate heterogeneity at the 5’-end of tRNA transcripts, which can arise from imprecise transcription initiation by T7 RNA polymerase(50,51). Further investigation will be required to determine how such leader sequences behave in other sequence contexts and whether additional optimization is possible.

In this study, we observed differences in processing efficiency among four IPEN arrangements (IPEN, PIEN, EIPN, and NIPE) (Figs. 3B and 3D), as well as among PIEN_v1, PIEN_v2, and L_PIEN_v2 (Fig. S3). To investigate whether these differences were associated with structural changes, we analyzed the predicted secondary structures using ViennaRNA (Fig. S7). The precursor RNAs of the EIPN and NIPE constructs, both of which showed lower HDV ribozyme (HDVR) self-cleavage efficiency, adopted relatively similar predicted structures both with (Fig. S7A) and without (Fig. S7B) the HDVR and terminator sequences. However, we have no direct evidence that this structural tendency accounts for the reduced cleavage efficiency. Likewise, no clear structural tendencies were identified among the predicted structures of PIEN_v1, PIEN_v2, and L_PIEN_v2 (Fig. S7C). Taken together, these results suggest that the mechanistic basis of cleavage efficiency remains poorly understood, and elucidating this mechanism represents an important direction for future research. Although tRNA array version 2 constructed in this study exhibited high translation performance as a simultaneous expression system based on unmodified tRNAs, there remains considerable room for further improvement. For example, the same optimization strategy as conducted for the PIEN group in this study can be applied to the other tRNA groups. In addition, because both HDVR and RNase P processing efficiencies are sequence dependent, systematic design of tRNA order within the array and of leader sequences may enable more flexible control of tRNA composition tailored to specific translation demands. In the longer term, customization of the PURE system using translation factors better suited for unmodified tRNAs(52) could further advance this platform toward a self-reproducible translation system with translation performance approaching that of native tRNAs.

## Acknowledgement

The authors declare no conflict of interest associated with this manuscript. This work was supported by JST, CREST Grant Number JPMJCR20S1, Japan, and Kakenhi Grant Numbers 22H05402, 24H01111, and 23KJ0815.

Ryota Miyachi: Conceptualization, Methodology, Data curation, Investigation, Writing - Original Draft. Anna Irie: Methodology, Investigation. Norikazu Ichihashi: Conceptualization, Supervision, Writing-Reviewing and Editing, Funding acquisition.

Supporting Information includes supplemental figures (Figures S1–S7) and tables (Tables S1–S6).

