## Supplemental figures and tables for "Enhanced tRNA array method version 2 for simultaneous in vitro synthesis of 21 tRNAs"

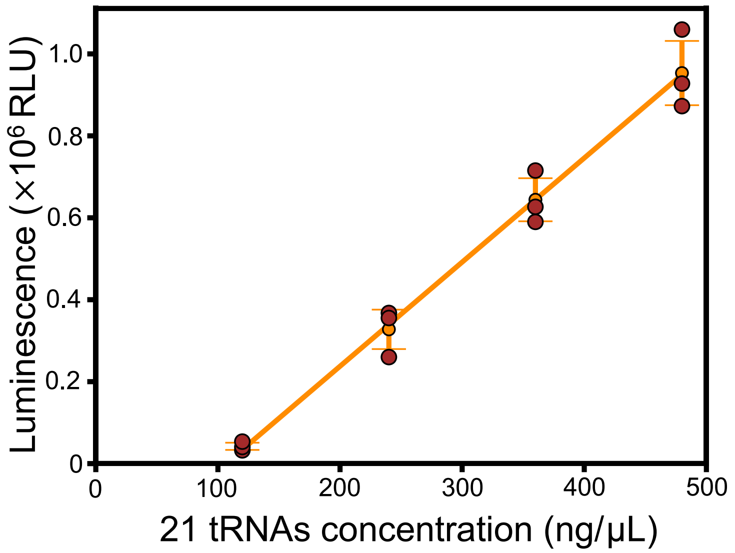


**Figure S1. Dependency of translation on 21 tRNA concentrations**

The 21 tRNAs mixture was added to the tRNA-free PURE system (tfPURE, composition A) together with a luciferase-encoding DNA template (1 nM) and T7 RNA polymerase (0.42 U/μL). Reactions were incubated at 30 °C for 16 h, and the luciferase activity was measured. Brown dots represent results from three independent experiments, orange dots indicate mean values, and error bars represent standard deviations. The orange line shows a linear regression fitted to the mean values.

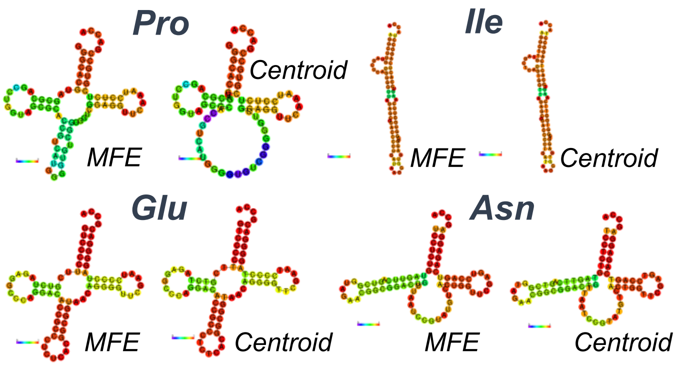


**Figure S2. Secondary structure prediction of tRNA$^{Pro}$, tRNA$^{Ile}$, tRNA$^{Glu}$, and tRNA$^{Asn}$ used in the previous tRNA array method**

Secondary structures of tRNA$^{Pro}$, tRNA$^{Ile}$, tRNA$^{Glu}$, and tRNA$^{Asn}$ encoded in the tRNA array were predicted using the RNAfold Web Server from the ViennaRNA website. The minimum free energy (MFE) structure represents the single secondary structure with the lowest predicted free energy, whereas the centroid structure represents the structure that best reflects the ensemble of probable structures based on base-pairing probabilities. Colors indicate the probability of base pairing from purple (low) to red (high).

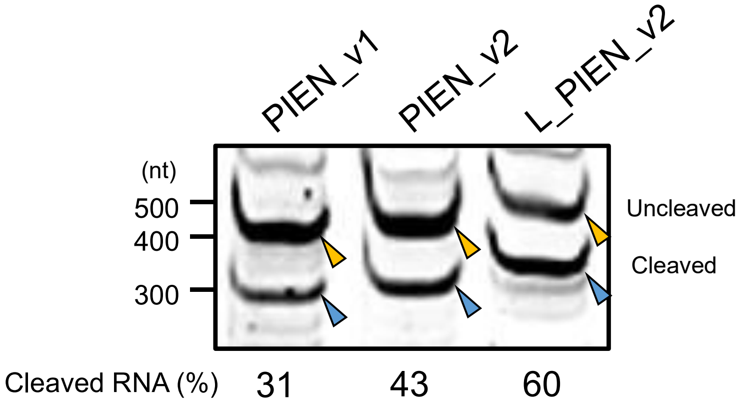


**Figure S3. Transcription and HDVR self-cleavage of each PIEN constructs**

Urea-PAGE analysis of transcripts generated from the previous construct (PIEN_v1), PIEN_v2, and L_PIEN_v2 templates, followed by SYBR green II staining. Predicted bands corresponding to the uncleaved transcript containing HDVR and the transcript after self-cleavage of HDVR are indicated by yellow and blue arrowheads, respectively.

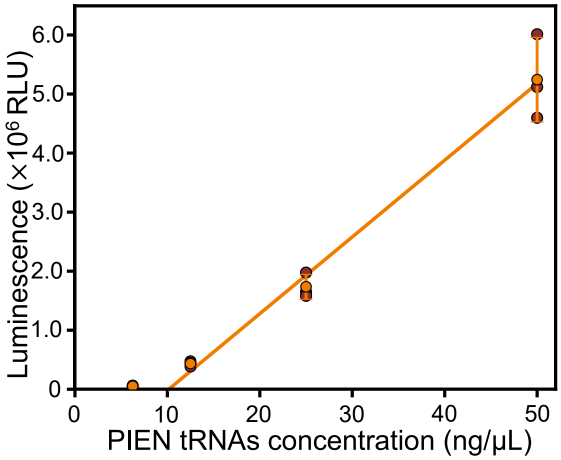


**Figure S4. Dependence of luciferase translation on PIEN tRNAs concentration**

The PIEN tRNAs mixture was added to the tRNA-free PURE system (tfPURE, composition A) together with a 17 tRNAs mixture (12 ng/μL each), luciferase-encoding DNA template (1 nM), and T7 RNA polymerase (0.42 U/μL). Reactions were incubated at 30 °C for 16 h, and the luciferase activity was measured. Dots represent results from three independent experiments, orange dots indicate mean values, and error bars represent standard deviations. The orange line indicates linear regression using the three higher concentrations (12.5, 25, and 50 ng/μL).

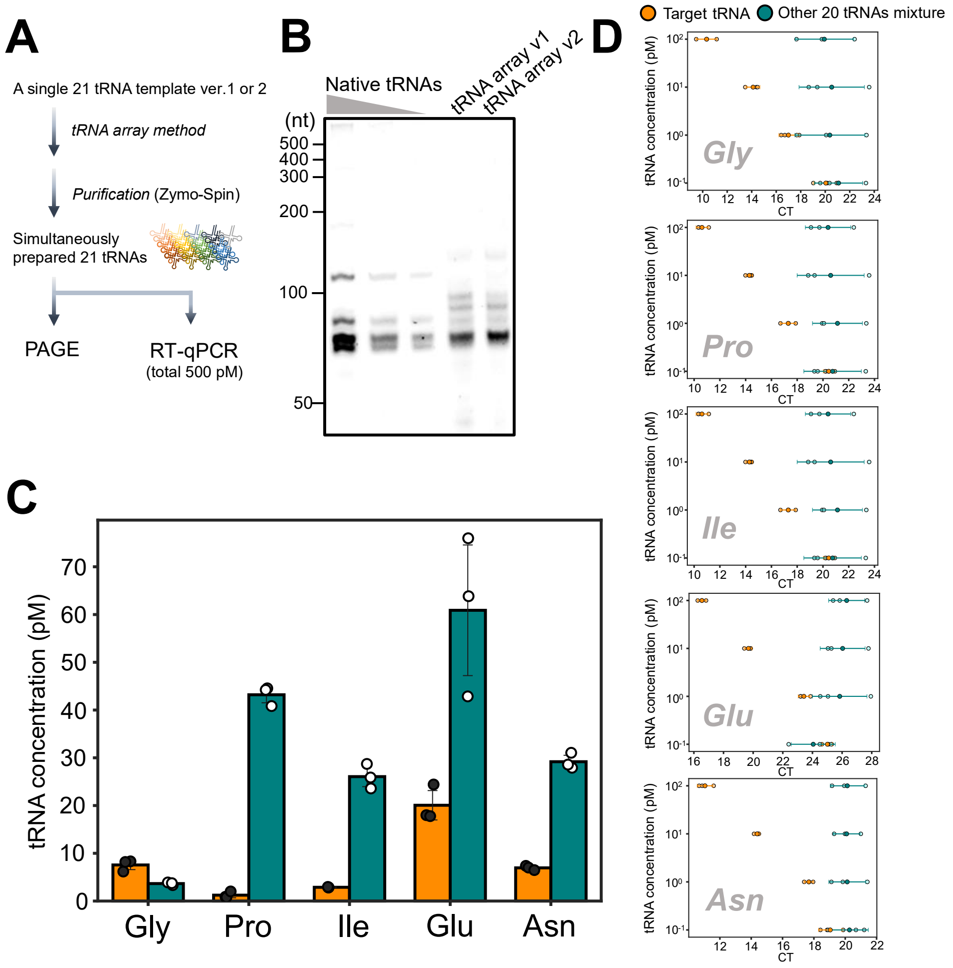


**Figure S5. Quantification of selected tRNA species in purified 21-tRNA mixtures by RT-qPCR.**

(A) Experimental workflow for RT-qPCR analysis of purified 21-tRNA mixtures prepared from tRNA array version 1 and version 2 constructs. After in vitro transcription and processing, mature-sized tRNAs were purified and subjected to Urea-PAGE analysis or RT-qPCR. (B) Urea-PAGE analysis of purified 21 tRNAs prepared using tRNA array version 1 or version 2. A mixture of native E. coli tRNAs (Roche) was used as a reference for the size. (C) Quantified abundances of selected tRNA species determined by RT-qPCR. The four PIEN-group tRNAs ($tRNA^{Pro}$, $tRNA^{Ile}$, $tRNA^{Glu}$, and $tRNA^{Asn}$) and $tRNA^{Gly}$ as a control were analyzed. Quantification was performed using standard curves generated from serial dilutions of tRNAs with known concentrations. Bars indicate mean values, and error bars indicate standard deviations. (D) Validation of primer specificity. Each primer set was tested against the corresponding target tRNA (orange) and the other 20 tRNAs mixture (green) to confirm selective amplification. For 20 tRNA mixture, each of the 20 tRNA concentrations is indicated on the vertical axis. Darker points indicate mean values, and lighter points indicate individual replicate measurements (N = 3). Error bars indicate standard deviations.

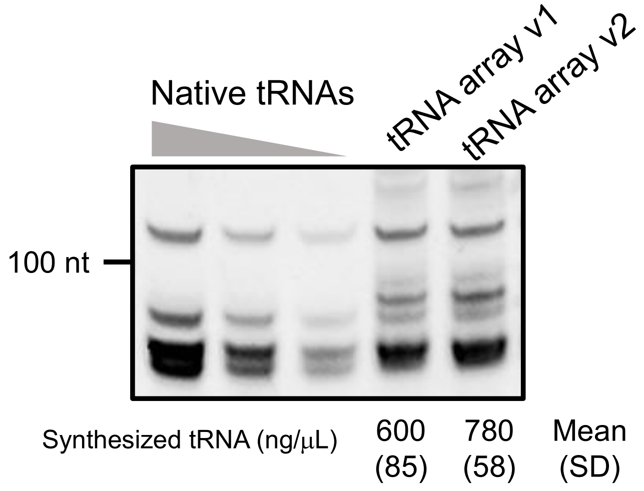


**Figure S6. Synthesis of 21 tRNAs in the tfPURE system using tRNA array version 1 or version 2**

Urea-PAGE analysis of the 21 tRNA products obtained after the translation-coupled reaction shown in Fig. 5E. A mixture of native E. coli tRNAs (Roche) was used as the reference for tRNA quantification. The band intensities corresponding to tRNAs were quantified using ImageJ by comparing to those of native tRNAs at known concentrations.

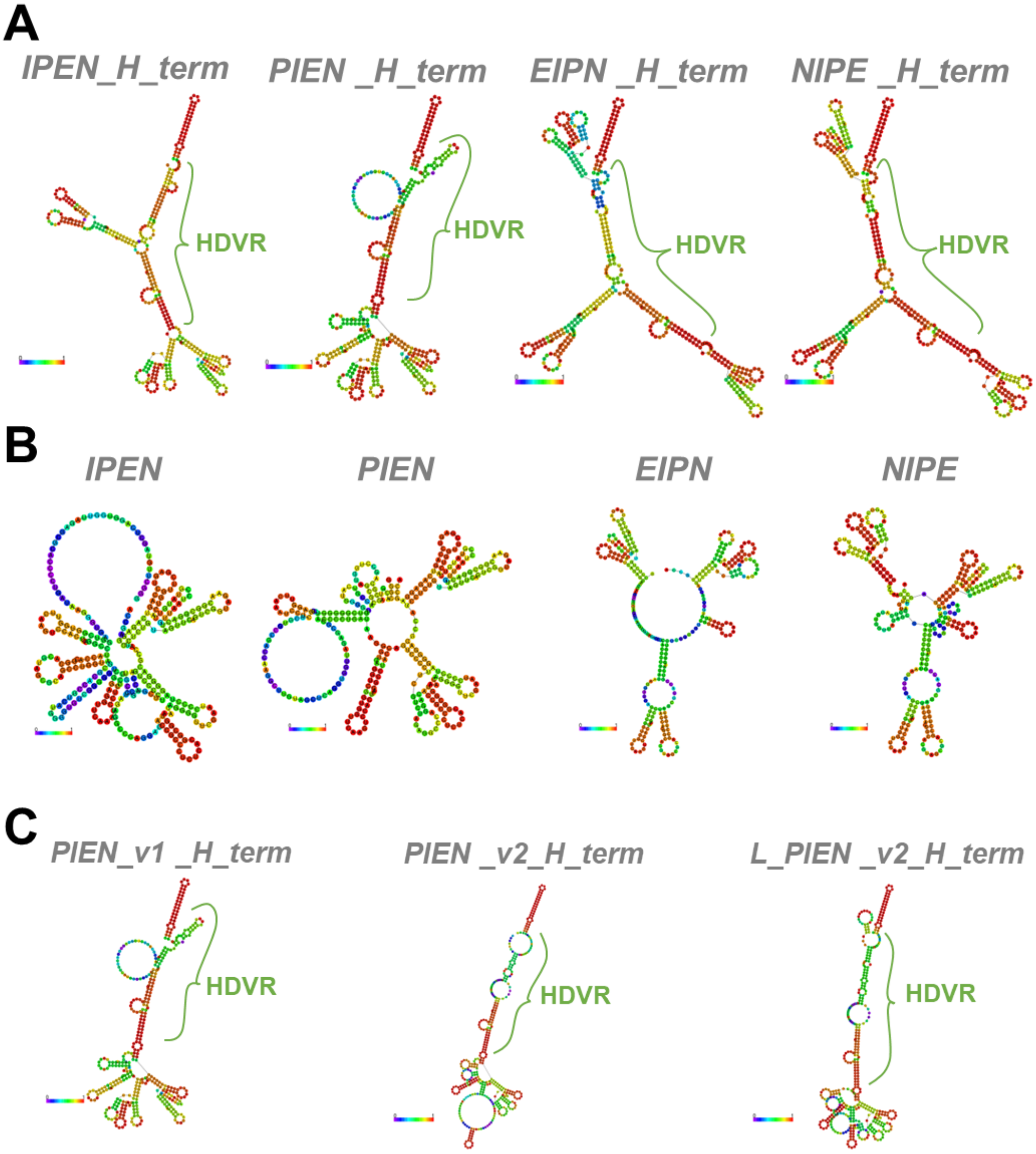


**Figure S7. Predicted secondary structures of precursor RNAs**

(A) Centroid secondary structures of the full-length precursor RNAs containing the tRNA array, HDV ribozyme (HDVR), and terminator sequence for the IPEN, PIEN, EIPN, and NIPE arrangements. The HDVR-containing region is indicated by green brackets. (B) Centroid structures of the corresponding tRNA-array regions without the HDVR and terminator sequences. (C) Centroid secondary structures of the full-length precursor RNAs containing the tRNA array, HDV ribozyme (HDVR), and terminator sequence for PIEN_v1, PIEN_v2, and L_PIEN_v2. The HDVR-containing region is indicated by green brackets. All structures were predicted using the RNAfold Web Server from the ViennaRNA website. Colors indicate the probability of base pairing from purple (low) to red (high).

**Table S1. Primer sets used in this study**

| Primer No. | Sequence |
|---|---|
| 1 | AATTCTAATACGACTCACTATAGGGCTTGTAGCTCAGATGGTTAGAGCGCACCCCTGATAAGGGTGAGGTCGGTGGTT CAAGTCCACTCAGGCCTACCA |
| 2 | AATTCTAATACGACTCACTATAGGGCTTGTAGCTCAGGTTGTTAGAGCGCACCCCTGATAAGGGTGAGGTCGGTGGTT CAAGTCCACTCAGGCCTACCA |
| 3 | AATTCTAATACGACTCACTATAGGGCTTGTAGCTCAGGTGGTTAGAGCGCACCCGTGATAAGGGTGAGGTCGGTGGTT CAAGTCCACTCAGGCCTACCA |
| 4 | AATTCTAATACGACTCACTATAGGGCTTGTAGCTCAGGTGGTTAGAGCGCACCCCAGATAAGGGTGAGGTCGGTGGTT CAAGTCCACTCAGGCCTACCA |
| 5 | AATTCTAATACGACTCACTATAGGGCTTGTAGCTCAGGTGGTTAGAGCGCACCCCTGATAAGGGTGAAGTCGGTGGTT CAAGTCCACTCAGGCCTACCA |
| 6 | TGGTAGGCCTGAGTGGACTTG |
| 7 | AATTCTAATACGACTCACTATAGCCTCTGTAGTTCAGTAGGTAGAACGGCGGACTGTTAATCCGTATGTCACTGGTTCG AGTCCAGTCAGAGGCGCCA |
| 8 | AATTCTAATACGACTCACTATAGCCTCTGTAGTTCAGTGGGTAGAACGGCGGACTGTTAATCCGTATGTCACTGGTTCG AGTCCAGTCAGAGGCGCCA |
| 9 | AATTCTAATACGACTCACTATAGCCTCTGTAGTTCAGTCAGTAGAACGGCGGACTGTTAATCCGTATGTCACTGGTTCG AGTCCAGTCAGAGGCGCCA |
| 10 | AATTCTAATACGACTCACTATAGCCTCTGTAGTTCAGTCGGTAGAACGGCGGAGTGTTAATCCGTATGTCACTGGTTCG AGTCCAGTCAGAGGCGCCA |
| 11 | AATTCTAATACGACTCACTATAGCCTCTGTAGTTCAGTCGGTAGAACGGCGGACTGTTAGTCCGTATGTCACTGGTTCG AGTCCAGTCAGAGGCGCCA |
| 12 | TGGCGCCTCTGACTGGACTCG |
| 13 | AATTCTAATACGACTCACTATAGGGCACGTAGCGCAGCCTGGTAGCGCACCGACATGGGGTGTCGGGGGTCGGAGGTT CAAATCCTCTCGTGCCGACCA |
| 14 | AATTCTAATACGACTCACTATAGGGCACGTAGCGCAGCCTGGTAGCGCACCGTCATGGGGTGACGGGGGTCGGAGGTT CAAATCCTCTCGTGCCGACCA |
| 15 | AATTCTAATACGACTCACTATACGGCACGTAGCGCAGCCTGGAAGCGCACCGTCATGGGGTGACGGGGGTCGGAGGT TCAAATCCTCTCGTGCCGACCA |
| 16 | AATTCTAATACGACTCACTATACGGCACGTAGCGCAGACTGGTAGCGCACCGTCATGGGGTGACGGGGGTCGGAGGTT CAAATCCTCTCGTGCCGACCA |
| 17 | AATTCTAATACGACTCACTATACGGCACGTAGCGCAGCCTGATAGCGCACCGTCATGGGGTGACGGGGGTCGGAGGTT CAAATCCTCTCGTGCCGACCA |
| 18 | TGGTCGGCACGAGAGGATTT |
| 19 | GAGATTAATACGACTCACTATAGGGCACGTAGCGCAGCCTGG |
| 20 | CGACTCACTATAGGGATAATACAACGGTTTC |
| 21 | TGAGCGGATAACAATTTCACACAGG |
| 22 | ATTGTTATCCGCTCAGAGGGCACAA |
| 23 | CCCTATAGTGAGTCGTATTAGAATTTGATCATG |
| 24 | AAGCGGAAGAGCGCCCAATACGC |
| 25 | GGCGATTAAGTTGGGTAACGCCAG |
| 26 | CCGGCTCGTATGTTGTGTGG |
| 27 | GGGGCTGCAGTAATACGACTCACTATA |
| 28 | AGGTGAAACTGACCGATAAG |
| 29 | AATAGCTCAGTTGGTAGAGCACGACCT |
| 30 | AGCGGGAAACGAGACTCGAACT |
| 31 | CCTGGTAGCGCACCGTCA |
| 32 | TCGGCACGAGAGGATTTGAACCTCC |
| 33 | CCTGGAAGCGCACCGTCA |
| 34 | TTGTAGCTCAGGTGGTTAGAGCGCACC |
| 35 | AGTGGACTTGAACCACCGACCTCACC |
| 36 | TTGTAGCTCAGATGGTTAGAGCGCACC |
| 37 | TTCGTCTAGAGGCCCAGGA |
| 38 | CCTGTTACCGCCGTGAGAGGG |

| | |
|---|---|
| 39 | TCCTCTGTAGTTCAGTCGGTAGAACGG |
| 40 | TGGACTCGAACCAGTGACATACGGATTA |
| 41 | GCCTCTGTAGTTCAGTGGGTAGAACGG |

### Table S2. Sequences of tRNA variants and primer sets for their preparation

| Primer sets | tRNA | tRNA seq |
|---|---|---|
| - | $tRNA^{Ile}$GAU_original | GGGCUUGUAGCUCAGGUGGUUAGAGCGCACCCCUGAUAAGG GUGAGGUCGGUGGUUCAAGUCCACUCAGGCCUACCA |
| 1,6 | $tRNA^{Ile}$GAU_G16A | GGGCUUGUAGCUCAGAUGGUUAGAGCGCACCCCUGAUAAGG GUGAGGUCGGUGGUUCAAGUCCACUCAGGCCUACCA |
| 2,6 | $tRNA^{Ile}$GAU_G18U | GGGCUUGUAGCUCAGGUUGUUAGAGCGCACCCCUGAUAAGG GUGAGGUCGGUGGUUCAAGUCCACUCAGGCCUACCA |
| 3,6 | $tRNA^{Ile}$GAU_C32G | GGGCUUGUAGCUCAGGUGGUUAGAGCGCACCCGUGAUAAGG GUGAGGUCGGUGGUUCAAGUCCACUCAGGCCUACCA |
| 4,6 | $tRNA^{Ile}$GAU_U33A | GGGCUUGUAGCUCAGGUGGUUAGAGCGCACCCCAGAUAAGG GUGAGGUCGGUGGUUCAAGUCCACUCAGGCCUACCA |
| 5,6 | $tRNA^{Ile}$GAU_G45A | GGGCUUGUAGCUCAGGUGGUUAGAGCGCACCCCUGAUAAGG GUGAAGUCGGUGGUUCAAGUCCACUCAGGCCUACCA |
| - | $tRNA^{Asn}$GUU_original | GCCUCUGUAGUUCAGUCGGUAGAACGGCGGACUGUUAAUCC GUAUGUCACUGGUUCGAGUCCAGUCAGAGGCGCCA |
| 7,12 | $tRNA^{Asn}$GUU_C17A | GCCUCUGUAGUUCAGUAGGUAGAACGGCGGACUGUUAAUCC GUAUGUCACUGGUUCGAGUCCAGUCAGAGGCGCCA |
| 8,12 | $tRNA^{Asn}$GUU_C17G | GCCUCUGUAGUUCAGUGGGUAGAACGGCGGACUGUUAAUCC GUAUGUCACUGGUUCGAGUCCAGUCAGAGGCGCCA |
| 9,12 | $tRNA^{Asn}$GUU_G18A | GCCUCUGUAGUUCAGUCAGUAGAACGGCGGACUGUUAAUCC GUAUGUCACUGGUUCGAGUCCAGUCAGAGGCGCCA |
| 10,12 | $tRNA^{Asn}$GUU_C32G | GCCUCUGUAGUUCAGUCGGUAGAACGGCGGAGUGUUAAUCC GUAUGUCACUGGUUCGAGUCCAGUCAGAGGCGCCA |
| 11,12 | $tRNA^{Asn}$GUU_A38G | GCCUCUGUAGUUCAGUCGGUAGAACGGCGGACUGUUAGUCC GUAUGUCACUGGUUCGAGUCCAGUCAGAGGCGCCA |
| - | $tRNA^{Pro}$GGG_original | GGGCACGUAGCGCAGCCUGGUAGCGCACCGUCAUGGGGUGU CGGGGGUCGGAGGUUCAAAUCCUCUCGUGCCGACCA |
| 13,18 | $tRNA^{Pro}$GGG_U30A | GGGCACGUAGCGCAGCCUGGUAGCGCACCGACAUGGGGUGU CGGGGGUCGGAGGUUCAAAUCCUCUCGUGCCGACCA |
| 14,18 | $tRNA^{Pro}$GGG_U40A | GGGCACGUAGCGCAGCCUGGUAGCGCACCGUCAUGGGGUGA CGGGGGUCGGAGGUUCAAAUCCUCUCGUGCCGACCA |
| 15,18,19 | $tRNA^{Pro}$GGG_U40A _U20A | GGGCACGUAGCGCAGCCUGGAAGCGCACCGUCAUGGGGUGA CGGGGGUCGGAGGUUCAAAUCCUCUCGUGCCCACCA |
| - | tRNAProGGG_G1C | CGGCACGUAGCGCAGCCUGGUAGCGCACCGUCAUGGGGUGU CGGGGGUCGGAGGUUCAAAUCCUCUCGUGCCGACCA |
| 16,18 | $tRNA^{Pro}$GGG_G1C_U40A_C 16A | CGGCACGUAGCGCAGACUGGUAGCGCACCGUCAUGGGGUGA CGGGGGUCGGAGGUUCAAAUCCUCUCGUGCCGACCA |
| 17,18 | $tRNA^{Pro}$GGG_G1C_U40A _G19A | CGGCACGUAGCGCAGCCUGAUAGCGCACCGUCAUGGGGUGA CGGGGGUCGGAGGUUCAAAUCCUCUCGUGCCGACCA |
| 15,18 | $tRNA^{Pro}$GGG_G1C_U40A _U20A | CGGCACGUAGCGCAGCCUGGAAGCGCACCGUCAUGGGGUGA CGGGGGUCGGAGGUUCAAAUCCUCUCGUGCCGACCA |

### Table S3. DNA sequences used in this study

| Template | Sequence |
|---|---|
| pUC-T7p-Fluc_21tRNA<br>*Fluc_gene | TCGCGCGTTTCGGTGATGACGGTGAAAACCTCTGACACATGCAGCTCCCGGAGACGGTCACAGCTTGTCTGTAAGCGGATGCCGGGAGCAGACAAGCCCGTCAGG<br>GCGCGTCAGCGGGTGTTGGCGGGTGTCGGGGCTGGCTTAACTATGCGGCATCAGAGCAGATTGTACTGAGAGTGCACCATATGCGGTGTGAAATACCGCACAGAT<br>GCGTAAGGAGAAAATACCGCATCAGGCGCCATTCGCCATTCAGGCTGCGCAACTGTTGGGAAGGGCGATCGGTGCGGGCCTCTTCGCTATTACGCCAGCTGGCGA<br>AAGGGGGATGTGCTGCAAGGCGATTAAGTTGGGTAACGCCAGGGTTTTCCCAGTCACGACGTTGTAAAACGACGGCCAGTGAATTCTAATACGACTCACTATAGG<br>GGAATTGTGAGCGGATAACAATTCCCCTCTAGAAATAATTTTGTTTAACTTTAAGAAGGAGATATACATATGGAGGACGCTAAGAACATTAAGAAGGGTCCTGCTC<br>CTTTTTACCCTCTGGAGGACGGTACTGCTGGTGAGCAGCTGCACAAGGCTATGAAGCGGTACGCTCTGGTTCCTGGTACTATTGCTTTTACTGACGCTCACATTGAG<br>GTTAACATTACTTACGCTGAGTACTTTGAGATGTCTGTTCGGCTGGCTGAGGCTATGAAGCGGTACGGTCTGAACACTAACCACCGGATTGTTGTTTGTTCTGAGAA<br>CTCTCTGCAGTTTTTTATGCCTGTTCTGGGTGCTCTGTTTATTGGTGTTGCTGTTGCTCCTGCTAACGACATTTACAACGAGCGGGAGCTGCTGAACTCTATGAACAT<br>TTCTCAGCCTACTGTTGTTTTTGTTTCTAAGAAGGGTCTGCAGAAGATTCTGAACGTTCAGAAGAAGCTGCCTATTATTCAGAAGATTATTATTATGGACTCTAAGA<br>CTGACTACCAGGGTTTTCAGTCTATGTACACTTTTGTTACTTCTCACCTGCCTCCTGGTTTTAACGAGTACGACTTTGTTCCTGAGTCTTTTGACCGGGACAAGACTA<br>TTGCTCTGATTATGAACTCTTCTGGTTCTACTGGTCTGCCTAAGGGTGTTGCTCTGCCTCACCGGACTGCTTGTGTTCGGTTTTCTCACGCTCGGGACCCTATTTTTGG<br>TAACCAGATTATTCCTGACACTGCTATTCTGTCTGTTGTTCCTTTTCACCACGGTTTTGGTATGTTTACTACTCTGGGTTACCTGATTTGTGGTTTTCGGGTTGTTCTG<br>ATGTACCGGTTTGAGGAGGAGCTGTTTCTGCGGTCTCTGCAGGACTACAAGATTCAGTCTGCTCTGCTGGTTCCTACTCTGTTTTCTTTTTTTGCTAAGTCTACTCTG<br>ATTGACAAGTACGACCTGTCTAACCTGCACGAGATTGCTTCTGGTGGTGCTCCTCTGTCTAAGGAGGTTGGTGAGGCTGTTGCTAAGCGGTTTCACCTGCCTGGTAT<br>TCGGCAGGGTTACGGTCTGACTGAGACTACTTCTGCTATTCTGATTACTCCTGAGGGTGACGACAAGCCTGGTGCTGTTGGTAAGGTTGTTCCTTTTTTTGAGGCTA<br>AGGTTGTTGACCTGGACACTGGTAAGACTCTGGGTGTTAACCAGCGGGGTGAGCTGTGTGTTCGGGGTCCTATGATTATGTCTGGTTACGTTAACAACCCTGAGGCT<br>ACTAACGCTCTGATTGACAAGGACGGTTGGCTGCACTCTGGTGACATTGCTTACTGGGACGAGGACGAGCACTTTTTTATTGTTGACCGGCTGAAGTCTCTGATTAA<br>GTACAAGGGTTACCAGGTTGCTCCTGCTGAGCTGGAGTCTATTCTGCTGCAGCACCCTAACATTTTTGACGCTGGTGTTGCTGGTCTGCCTGACGACGACGCTGGTG<br>AGCTGCCTGCTGCTGTTGTTGTTCTGGAGCACGGTAAGACTATGACTGAGAAGGAGATTGTTGACTACGTTGCTTCTCAGGTTACTACTGCTAAGAAGCTGCGGGGT<br>GGTGTTGTTTTTGTTGACGAGGTTCCTAAGGGTCTGACTGGTAAGCTGGACGCTCGGAAGATTCGGGAGATTCTGATTAAGGCTAAGAAGGGTGGTAAGTCTAAGC<br>TGTAAAAGCTTGGCGTAATCATGGTCATAGCTGTTTCCTGTGTGAAATTGTTATCCGCTCACAATTCCACACAACATACGAGCCGGAAGCATAAAGTGTAAAGCCT<br>GGGGTGCCTAATGAGTGAGCTAACTCACATTAATTGCGTTGCGCTCACTGCCCGCTTTCCAGTCGGGAAACCTGTCGTGCCAGCTGCATTAATGAATCGGCCAACG<br>CGCGGGGAGAGGCGGTTTGCGTATTGGGCGCTCTTCCGCTTCCTCGCTCACTGACTCGCTGCGCTCGGTCGTTCGGCTGCGGCGAGCGGTATCAGCTCACTCAAAGG<br>CGGTAATACGGTTATCCACAGAATCAGGGGATAACGCAGGAAAGAACATGTGAGCAAAAGGCCAGCAAAAGGCCAGGAACCGTAAAAAGGCCGCGTTGCTGGCG<br>TTTTTCCATAGGCTCCGCCCCCCTGACGAGCATCACAAAAATCGACGCTCAAGTCAGAGGTGGCGAAACCCGACAGGACTATAAAGATACCAGGCGTTTCCCCCTG<br>GAAGCTCCCTCGTGCGCTCTCCTGTTCCGACCCTGCCGCTTACCGGATACCTGTCCGCCTTTCTCCCTTCGGGAAGCGTGGCGCTTTCTCATAGCTCACGCTGTAGGT<br>ATCTCAGTTCGGTGTAGGTCGTTCGCTCCAAGCTGGGCTGTGTGCACGAACCCCCCGTTCAGCCCGACCGCTGCGCCTTATCCGGTAACTATCGTCTTGAGTCCAAC<br>CCGGTAAGACACGACTTATCGCCACTGGCAGCAGCCACTGGTAACAGGATTAGCAGAGCGAGGTATGTAGGCGGTGCTACAGAGTTCTTGAAGTGGTGGCCTAAC<br>TACGGCTACACTAGAAGAACAGTATTTGGTATCTGCGCTCTGCTGAAGCCAGTTACCTTCGGAAAAAGAGTTGGTAGCTCTTGATCCGGCAAACAAACCACCGCTG<br>GTAGCGGTGGTTTTTTTGTTTGCAAGCAGCAGATTACGCGCAGAAAAAAAGGATCTCAAGAAGATCCTTTGATCTTTTCTACGGGGTCTGACGCTCAGTGGAACGA<br>AAACTCACGTTAAGGGATTTTGGTCATGAGATTATCAAAAAGGATCTTCACCTAGATCCTTTTAAATTAAAAATGAAGTTTTAAATCAATCTAAAGTATATATGAGT<br>AAACTTGGTCTGACAGTTACCAATGCTTAATCAGTGAGGCACCTATCTCAGCGATCTGTCTATTTCGTTCATCCATAGTTGCCTGACTCCCCGTCGTGTAGATAACTA<br>CGATACGGGAGGGCTTACCATCTGGCCCCAGTGCTGCAATGATACCGCGAGACCCACGCTCACCGGCTCCAGATTTATCAGCAATAAACCAGCCAGCCGGAAGGG<br>CCGAGCGCAGAAGTGGTCCTGCAACTTTATCCGCCTCCATCCAGTCTATTAATTGTTGCCGGGAAGCTAGAGTAAGTAGTTCGCCAGTTAATAGTTTGCGCAACGTT<br>GTTGCCATTGCTACAGGCATCGTGGTGTCACGCTCGTCGTTTGGTATGGCTTCATTCAGCTCCGGTTCCCAACGATCAAGGCGAGTTACATGATCCCCCATGTTGTG<br>CAAAAAAGCGGTTAGCTCCTTCGGTCCTCCGATCGTTGTCAGAAGTAAGTTGGCCGCAGTGTTATCACTCATGGTTATGGCAGCACTGCATAATTCTCTTACTGTCA<br>TGCCATCCGTAAGATGCTTTTCTGTGACTGGTGAGTACTCAACCAAGTCATTCTGAGAATAGTGTATGCGGCGACCGAGTTGCTCTTGCCCGGCGTCAATACGGGAT<br>AATACCGCGCCACATAGCAGAACTTTAAAAGTGCTCATCATTGGAAAACGTTCTTCGGGGCGAAAACTCTCAAGGATCTTACCGCTGTTGAGATCCAGTTCGATGT<br>AACCCACTCGTGCACCCAACTGATCTTCAGCATCTTTTACTTTCACCAGCGTTTCTGGGTGAGCAAAAACAGGAAGGCAAAATGCCGCAAAAAAGGGAATAAGGG<br>CGACACGGAAATGTTGAATACTCATACTCTTCCTTTTTCAATATTATTGAAGCATTTATCAGGGTTATTGTCTCATGAGCGGATACATATTTGAATGTATTTAGAAAA<br>ATAAACAAATAGGGGTTCCGCGCACATTTCCCCGAAAAGTGCCACCTGACGTCTAAGAAACCATTATTATCATGACATTAACCTATAAAAATAGGCGTATCACGAG<br>GCCCTTTCGTC |
| sfGFP_fragment_21tRNA<br>*sfGFP gene | GGCGATTAAGTTGGGTAACGCCAGGGTTTTCCCAGTCACGACGTTGTAAAACGACGGCCAGTGAATTCTAATACGACTCACTATAGGGGAATTGTGAGCGGATAAC<br>AATTCCCCTCTAGAAATAATTTTGTTTAACTTTAAGAAGGAGATATACATATGTCTAAGGGTGAGGAGCTGTTTACTGGTGTTGTTCCTATTCTGGTTGAGCTGGAC<br>GGTGACGTTAACGGTCACAAGTTTTCTGTTCGGGGTGAGGGTGAGGGTGACGCTACTAACGGTAAGCTGACTCTGAAGTTTATTTGTACTACTGGTAAGCTGCCTGT<br>TCCTTGGCCTACTCTGGTTACTACTCTGACTTACGGTGTTCAGTGTTTTTCTCGGTACCCTGACCACATGAAGCGGCACGACTTTTTTAAGTCTGCTATGCCTGAGGG<br>TTACGTTCAGGAGCGGACTATTTCTTTTAAGGACGACGGTACTTACAAGACTCGGGCTGAGGTTAAGTTTGAGGGTGACACTCTGGTTAACCGGATTGAGCTGAAG<br>GGTATTGACTTTAAGGAGGACGGTAACATTCTGGGTCACAAGCTGGAGTACAACTTTAACTCTCACAACGTTTACATTACTGCTGACAAGCAGAAGAACGGTATTA<br>AGGCTAACTTTAAGATTCGGCACAACGTTGAGGACGGTTCTGTTCAGCTGGCTGACCACTACCAGCAGAACACTCCTATTGGTGACGGTCCTGTTCTGCTGCCTGAC<br>AACCACTACCTGTCTACTCAGTCTGTTCTGTCTAAGGACCCTAACGAGAAGCGGGACCACATGGTTCTGCTGGAGTTTGTTACTGCTGCTGGTATTACTCACGGTAT<br>GGACGAGCTGTACAAGGGTTCTGTTTCTGGTTGGCGGCTGTTTAAGAAGATTTCTTGAAAAAGCTTGGCGTAATCATGGTCATAGCTGTTTCCTGTGTGAAATTGTT<br>ATCCGCTCACAATTCCACACAACATACGAGCCGG |
| GUS_21tRNA<br>*GUS gene | GGCGATTAAGTTGGGTAACGCCAGGGTTTTCCCAGTCACGACGTTGTAAAACGACGGCCAGTGAATTCTAATACGACTCACTATAGGGGAATTGTGAGCGGATAAC<br>AATTCCCCTCTAGAAATAATTTTGTTTAACTTTAAGAAGGAGATATACATATGGCTTCTATGACTGGTGGTCAGCAGATGGGTCGGGGTTCTGAGTTTGAGGCTCTG<br>CGGGCTCTGCGGCCTGTTGAGACTCCTACTCGGGAGATTAAGAAGCTGGACGGTCTGTGGGCTTTTTCTCTGGACCGGGAGAACTGTGGTATTGACCAGCGGTGGT<br>GGGAGTCTGCTCTGCAGGAGTCTCGGGCTATTGCTGTTCCTGGTTCTTTTAACGACCAGTTTGCTGACGCTGACATTCGGAACTACGCTGGTAACGTTTGGTACCAG<br>CGGGAGGTTTTTATTCCTAAGGGTTGGGCTGGTCAGCGGATTGTTCTGCGGTTTGACGCTGTTACTCACTACGGTAAGGTTTGGGTTAACAACCAGGAGGTTATGGA<br>GCACCAGGGTGGTTACACTCCTTTTGAGGCTGACGTTACTCCTTACGTTATTGCTGGTAAGTCTGTTCGGATTACTGTTTGTGTTAACAACGAGCTGAACTGGCAGA<br>CTATTCCTCCTGGTATGGTTATTACTGACGAGAACGGTAAGAAGAAGCAGTCTTACTTTCACGACTTTTTTAACTACGCTGGTATTCACCGGTCTGTTATGCTGTACA<br>CTACTCCTAACACTTGGGTTGACGACATTACTGTTGTTACTCACGTTGCTCAGGACTGTAACCACGCTTCTGTTGACTGGCAGGTTGTTGCTAACGGTGACGTTTCTG<br>TTGAGCTGCGGGACGCTGACCAGCAGGTTGTTGCTACTGGTCAGGGTACTTCTGGTACTCTGCAGGTTGTTAACCCTCACCTGTGGCAGCCTGGTGAGGGTTACCTG<br>TACGAGCTGTGTGTTACTGCTAAGTCTCAGACTGAGTGTGACATTTACCCTCTGCGGGTTGGTATTCGGTCTGTTGCTGTTAAGGGTGAGCAGTTTCTGATTAACCA<br>CAAGCCTTTTTACTTTACTGGTTTTGGTCGGCACGAGGACGCTGACCTGCGGGGTAAGGGTTTTGACAACGTTCTGATGGTTCACGACCACGCTCTGATGGACTGGA<br>TTGGTGCTAACTCTTACCGGACTTCTCACTACCCTTACGCTGAGGAGATGCTGGACTGGGCTGACGAGCACGGTATTGTTGTTATTGACGAGACTGCTGCTGTTGGT<br>TTTAACCTGTCTCTGGGTATTGGTTTTGAGGCTGGTAACAAGCCTAAGGAGCTGTACTCTGAGGAGGCTGTTAACGGTGAGACTCAGCAGGCTCACCTGCAGGCTA<br>TTAAGGAGCTGATTGCTCGGGACAAGAACCACCCTTCTGTTGTTATGTGGTCTATTGCTAACGAGCCTGACACTCGGCCTCAGGGTGCTCGGGAGTACTTTGCTCCT<br>CTGGCTGAGGCTACTCGGAAGCTGGACCCTACTCGGCCTATTACTTGTGTTAACGTTATGTTTTGTGACGCTCACACTGACACTATTTCTGACCTGTTTGACGTTCTG<br>TGTCTGAACCGGTACTACGGTTGGTACGTTCAGTCTGGTGACCTGGAGACTGCTGAGAAGGTTCTGGAGAAGGAGCTGCTGGCTTGGCAGGAGAAGCTGCACCAGC<br>CTATTATTATTACTGAGTACGGTGTTGACACTCTGGCTGGTCTGCACTCTATGTACACTGACATGTGGTCTGAGGAGTACCAGTGTGCTTGGCTGGACATGTACCAC<br>CGGGTTTTTGACCGGGTTTCTGCTGTTGTTGGTGAGCAGGTTTGGAACTTTGCTGACTTTGCTACTTCTCAGGGTATTCTGCGGGTTGGTGGTAACAAGAAGGGTATT<br>TTTACTCGGGACCGGAAGCCTAAGTCTGCTGCTTTTCTGCTGCAGAAGCGGTGGACTGGTATGAACTTTGGTGAGAAGCCTCAGCAGGGTGGTAAGCAGGGTCTGT<br>GTGGTCGGAAGCTGGCTGCTGCTCTGGAGGTTTCTGGTTGGCGGCTGTTTAAGAAGATTTCTTGAAAAAGCTTGGCGTAATCATGGTCATAGCTGTTTCCTGTGTGA<br>AATTGTTATCCGCTCACAATTCCACACAACATACGAGCCGG |
| L_PIEN_v2_fragment<br>*Leader<br>*PIEN_v2 | AAGCGGAAGAGCGCCCAATACGCAAACCGCCTCTCCCCGCGCGTTGGCCGATTCATTAATGCAGCCCGGGAGATCTCGATCCCGCGAAAAATTCTAATACGACTCA<br>CTATAGGGATAATACAACGGTTTCCCATATATCGGCACGTAGCGCAGCCTGGAAGCGCACCGTCATGGGGTGACGGGGGTCGGAGGTTCAAATCCTCTCGTGCCGA<br>CCAGGGCTTGTAGCTCAGATGGTTAGAGCGCACCCCTGATAAGGGTGAGGTCGGTGGTTCAAGTCCACTCAGGCCTACCAGTCCCCTTCGTCTAGAGGCCCAGGAC<br>ACCGCCCTCTCACGGCGGTAACAGGGGTTCGAATCCCCTAGGGGACGCCAGCCTCTGTAGTTCAGTGGGTAGAACGGCGGACTGTTAATCCGTATGTCACTGGTTC<br>GAGTCCAGTCAGAGGCGCCAGGGTCGGCATGGCATCTCCACCTCCTCGCGGTCCGACCTGGGCTACTTCGGTAGGCTAAGGGAGAAGAGATAACAGATACTTCGG<br>TATCTGTTATCTGTTTTTTTCAACAGATAGCCGCGTTCGCGCGGCTATCTGTTTTTTTGCTGAAAGGAGGAACTATATCCGGATAACCTGGCAGGCTAGGTGGAG<br>GCTCAGTGATGATAAGTCTGCGATGGTGGATGCATGTGTCATGGTCATAGCTGTTTCCTGTGTGAAATTGTTATCCGCTCA |
| pTwist_21tRNAs<br>*GARCQ group<br>*LWY group | AAGCGGAAGAGCGCCCAATACGCAAACCGCCTCTCCCCGCGCGTTGGCCGATTCATTAATGCAGCCCGGGAGATCTCGATCCCGCGAAAAATTCTAATACGACTCA<br>CTATAGCGGGAATAGCTCAGTTGGTAGAGCACGACCTTGCCAAGGTCGGGGTCGCGAGTTCGAGTCTCGTTTCCCGCTCCAGGGGCTATAGCTCAGCTGGGAGAGC<br>GCTTGCATGGCATGCAAGAGGTCAGCGGTTCGATCCCGCTTAGCTCCACCAGCGCCCGTAGCTCAGCTGGATAGAGCGCTGCCCTCCGGAGGCAGAGGTCTCAGGT<br>TCGAATCCTGTCGGGCGCGCCAGGCGCGTTAACAAAGCGGTTATGTAGCGGATTGCAAATCCGTCTAGTCCGGTTCGACTCCGGAACGCGCCTCCATGGGGTATCG<br>CCAAGCGGTAAGGCACCGGATTCTGATTCCGGCATTCCGAGGTTCGAATCCTCGTACCCCAGCCAGGGTCGGCATGGCATCTCCACCTCCTCGCGGTCCGACCTGG<br>GCTACTTCGGTAGGCTAAGGGAGAAGAGATAACAGATACTTCGGTATCTGTTATCTGTTTTTTTCAACAGATAGCCGCGTTCGCGCGGCTATCTGTTTTTTTAGGC<br>TAGGTGGAGGCTCAGTGATGCCGTACTGGCTAAAACCTACGGCGGTGCTGCTCGCGCATTCGACATTCTAATACGACTCACTATAGCGAAGGTGGCGGAATTGGTA<br>GACGCGCTAGCTTCAGGTGTTAGTGTCCTTACGGACGTGGGGGTTCAAGTCCCCCCCTCGCACCAAGGGGCGTAGTTCAATTGGTAGAGCACCGGTCTCCAAAAC<br>CGGGTGTTGGGAGTTCGAGTCTCTCCGCCCCTGCCAGGTGGGGTTCCCGAGCGGCCAAAGGGAGCAGACTGTAAATCTGCCGTCACAGACTTCGAAGGTTCGAATC |

| | |
|---|---|
| *DfMHKV group<br>*SmMFTgroup<br>*PIEN group | CTTCCCCCACCACCAGGGTCGGCATGGCATCTCCACCTCCTCGCGGTCCGACCTGGGCTACTTCGGTAGGCTAAGGGAGAAGAGATAACAGATACTTCGGTATCTG<br>TTATCTGTTTTTTTTCAACAGATAGCCGCGTTCGCGCGGCTATCTGTTTTTTTTAGGCTAGGTGGAGGCTCAGTGATGAGATCGATAACGCGCCGGAAGAAAAAGCT<br>CGTGGTATCACCATCAAATTCTAATACGACTCACTATAGGAGCGGTAGTTCAGTCGGTTAGAATACCTGCCTGTCACGCAGGGGGTCGCGGGTTCGAGTCCCGTCC<br>GTTCCGCCACGCGGGGTGGAGCAGCCTGGTAGCTCGTCGGGCTCATAACCCGAAGATCGTCGGTTCAAATCCGGCCCCCGCAACCAGGTGGCTATAGCTCAGTTGG<br>TAGAGCCCTGGATTGTGATTCCAGTTGTCGTGGGTTCGAATCCCATTAGCCACCCCAGGGTCGTTAGCTCAGTTGGTAGAGCAGTTGACTCTTAATCAATTGGTCGC<br>AGGTTCGAATCCTGCACGACCCACCAGCGTCCGTAGCTCAGTTGGTTAGAGCACCACCTTGACATGGTGGGGGTCGGTGGTTCGAGTCCACTCGGACGCACCAGGG<br>TCGGCATGGCATCTCCACCTCCTCGCGGTCCGACCTGGGCTACTTCGGTAGGCTAAGGGAGAAGAGATAACAGATACTTCGGTATCTGTTATCTGTTTTTTTTCAAC<br>AGATAGCCGCGTTCGCGCGGCTATCTGTTTTTTTTAGGCTAGGTGGAGGCTCAGTGATGACACTTCTCACGTTGAATACGACACCCCGACCCGTCACTACGCACAAT<br>TCTAATACGACTCACTATAGGTGAGGTGTCCGAGTGGCTGAAGGAGCACGCCTGGAAAGTGTGTATACGGCAACGTATCGGGGGTTCGAATCCCCCCCTCACCGCC<br>AGGCTACGTAGCTCAGTTGGTTAGAGCACATCACTCATAATGATGGGGTCACAGGTTCGAATCCCGTCGTAGCCACCAGCCCGGATAGCTCAGTCGGTAGAGCAGG<br>GGATTGAAAATCCCCGTGTCCTTGGTTCGATTCCGAGTCCGGGCACCAGCTGATATGGCTCAGTTGGTAGAGCGCACCCTTGGTAAGGGTGAGGTCCCCAGTTCGA<br>CTCTGGGTATCAGCACCAGGGTCGGCATGGCATCTCCACCTCCTCGCGGTCCGACCTGGGCTACTTCGGTAGGCTAAGGGAGAAGAGATAACAGATACTTCGGTAT<br>CTGTTATCTGTTTTTTTTCAACAGATAGCCGCGTTCGCGCGGCTATCTGTTTTTTTTAGGCTAGGTGGAGGCTCAGTGATGACGTAGACTGCCCGGGGCACGCCGAC<br>TATGTTAAAAACATGATCAAATTCTAATACGACTCACTATAGGGCACGTAGCGCAGCCTGGTAGCGCACCGTCATGGGGTGTCGGGGGTCGGAGGTTCAAATCCTC<br>TCGTGCCGACCAAGGCTTGTAGCTCAGGTGGTTAGAGCGCACCCCTGATAAGGGTGAGGTCGGTGGTTCAAGTCCACTCAGGCCTACCAGTCCCCTTCGTCTAGAG<br>GCCCAGGACACCGCCCTCTCACGGCGGTAACAGGGGTTCGAATCCCCTAGGGGACGCCATCCTCTGTAGTTCAGTCGGTAGAACGGCGGACTGTTAATCCGTATGT<br>CACTGGTTCGAGTCCAGTCAGAGGAGCCAGGGTCGGCATGGCATCTCCACCTCCTCGCGGTCCGACCTGGGCTACTTCGGTAGGCTAAGGGAGAAGAGATAACAG<br>ATACTTCGGTATCTGTTATCTGTTTTTTTTCAACAGATAGCCGCGTTCGCGCGGCTATCTGTTTTTTTTGCTGAAAGGAGGAACTATATCCGGATAACCTGGCAGGCT<br>AGGTGGAGGCTCAGTGATGATAAGTCTGCGATGGTGGATGCATGTGTCATGGTCATAGCTGTTTCCTGTGTGAAATTGTTATCCGCTCAGAGGGCACAATCCTATTC<br>CGCGCTATCCGACAATCTCCAAGACATTAGGTGGAGTTCAGTTCGGCGTATGGCATATGTCGCTGGAAAGAACATGTGAGCAAAAGGCCAGCAAAAGGCCAGGAA<br>CCGTAAAAAGGCCGCGTTGCTGGCGTTTTTCCATAGGCTCCGCCCCCCTGACGAGCATCACAAAAATCGACGCTCAAGTCAGAGGTGGCGAAACCCGACAGGACT<br>ATAAAGATACCAGGCGTTTCCCCCTGGAAGCTCCCTCGTGCGCTCTCCTGTTCCGACCCTGCCGCTTACCGGATACCTGTCCGCCTTTCTCCCTTCGGGAAGCGTGG<br>CGCTTTCTCATAGCTCACGCTGTAGGTATCTCAGTTCGGTGTAGGTCGTTCGCTCCAAGCTGGGCTGTGTGCACGAACCCCCCGTTCAGCCCGACCGCTGCGCCTTA<br>TCCGGTAACTATCGTCTTGAGTCCAACCCGGTAAGACACGACTTATCGCCACTGGCAGCAGCCACTGGTAACAGGATTAGCAGAGCGAGGTATGTAGGCGGTGCTA<br>CAGAGTTCTTGAAGTGGTGGCCTAACTACGGCTACACTAGAAGAACAGTATTTGGTATCTGCGCTCTGCTGAAGCCAGTTACCTTCGGAAAAAGAGTTGGTAGCTC<br>TTGATCCGGCAAACAAACCACCGCTGGTAGCGGTGGTTTTTTTGTTTGCAAGCAGCAGATTACGCGCAGAAAAAAAGGATCTCAAGAAGATCCTTTGATCTTTTCT<br>ACGGGGTCTGACGCTCTATTCAACAAAGCCGCCGTCCCGTCAAGTCAGCGTAAATGGGTAGGGGGCTTCAAATCGTCCTCGTGATACCAATTCGGAGCCTGCTTTTT<br>TGTACAAACTTGTTGATAATGGCAATTCAAGGATCTTCACCTAGATCCTTTTAAATTAAAAATGAAGTTTTAAATCAATCTAAAGTATATATGAGTAAACTTGGTCT<br>GACAGTTACCAATGCTTAATCAGTGAGGCACCTATCTCAGCGATCTGTCTATTTCGTTCATCCATAGTTGCCTGACTCCCCGTCGTGTAGATAACTACGATACGGGA<br>GGGCTTACCATCTGGCCCCAGTGCTGCAATGATACCGCGAGAGCCACGCTCACCGGCTCCAGATTTATCAGCAATAAACCAGCCAGCCGGAAGGGCCGAGCGCAG<br>AAGTGGTCCTGCAACTTTATCCGCCTCCATCCAGTCTATTAATTGTTGCCGGGAAGCTAGAGTAAGTAGTTCGCCAGTTAATAGTTTGCGCAACGTTGTTGCCATTG<br>CTACAGGCATCGTGGTGTCACGCTCGTCGTTTGGTATGGCTTCATTCAGCTCCGGTTCCCAACGATCAAGGCGAGTTACATGATCCCCCATGTTGTGCAAAAAAGCG<br>GTTAGCTCCTTCGGTCCTCCGATCGTTGTCAGAAGTAAGTTGGCCGCAGTGTTATCACTCATGGTTATGGCAGCACTGCATAATTCTCTTACTGTCATGCCATCCGTA<br>AGATGCTTTTCTGTGACTGGTGAGTACTCAACCAAGTCATTCTGAGAATAGTGTATGCGGCGACCGAGTTGCTCTTGCCCGGCGTCAATACGGGATAATACCGCGC<br>CACATAGCAGAACTTTAAAAGTGCTCATCATTGGAAAACGTTCTTCGGGGCGAAAACTCTCAAGGATCTTACCGCTGTTGAGATCCAGTTCGATGTAACCCACTCG<br>TGCACCCAACTGATCTTCAGCATCTTTTACTTTCACCAGCGTTTCTGGGTGAGCAAAAACAGGAAGGCAAAATGCCGCAAAAAAGGGAATAAGGGCGACACGGAA<br>ATGTTGAATACTCATACTCTTCCTTTTTCAATATTATTGAAGCATTTATCAGGGTTATTGTCTCATGAGCGGATACATATTTGAATGTATTTAGAAAAATAAACAAAT<br>AGGGGTTCCGCGCACATTTCCCCGAAAAGTGCCAGATACCTGAAACAAAACCCATCGTACGGCCAAGGAAGTCTCCAATAACTGTGATCCACCACAAGCGCCAGG<br>GTTTTCCCAGTCACGACGTTGTAAAACGACGGCCAGTCATGCATAATCCGCACGCATCTGGAATAAGGAAGTGCCATTCCGCCTGACCT |
| pTwist_21tRNAs_v2<br>*GARCQ group<br>*LWY group<br>*DfMHKV group<br>*SmMFTgroup<br>*L_PIEN_v2 group | AAGCGGAAGAGCGCCCAATACGCAAACCGCCTCTCCCCGCGCGTTGGCCGATTCATTAATGCAGCCCGGGAGATCTCGATCCCGCGAAAAATTCTAATACGACTCA<br>CTATAGCGGGAATAGCTCAGTTGGTAGAGCACGACCTTGCCAAGGTCGGGGTCGCGAGTTCGAGTCTCGTTTCCCGCTCCAGGGGCTATAGCTCAGCTGGGAGAGC<br>GCTTGCATGGCATGCAAGAGGTCAGCGGTTCGATCCCGCTTAGCTCCACCAGCGCCCGTAGCTCAGCTGGATAGAGCGCTGCCCTCCGGAGGCAGAGGTCTCAGGT<br>TCGAATCCTGTCGGGCGCGCCAGGCGCGTTAACAAAGCGGTTATGTAGCGGATTGCAAATCCGTCTAGTCCGGTTCGACTCCGGAACGCGCCTCCATGGGGTATCG<br>CCAAGCGGTAAGGCACCGGATTCTGATTCCGGCATTCCGAGGTTCGAATCCTCGTACCCCAGCCAGGGTCGGCATGGCATCTCCACCTCCTCGCGGTCCGACCTGG<br>GCTACTTCGGTAGGCTAAGGGAGAAGAGATAACAGATACTTCGGTATCTGTTATCTGTTTTTTTTCAACAGATAGCCGCGTTCGCGCGGCTATCTGTTTTTTTTAGGC<br>TAGGTGGAGGCTCAGTGATGCCGTACTGGCTAAAACCTACGGCGGTGCTGCTCGCGCATTCGACATTCTAATACGACTCACTATAGCGAAGGTGGCGGAATTGGTA<br>GACGCGCTAGCTTCAGGTGTTAGTGTCCTTACGGACGTGGGGGTTCAAGTCCCCCCCCTCGCACCAAGGGGCGTAGTTCAATTGGTAGAGCACCGGTCTCCAAAAC<br>CGGGTGTTGGGAGTTCGAGTCTCTCCGCCCCTGCCAGGTGGGGTTCCCGAGCGGCCAAAGGGAGCAGACTGTAAATCTGCCGTCACAGACTTCGAAGGTTCGAATC<br>CTTCCCCCACCACCAGGGTCGGCATGGCATCTCCACCTCCTCGCGGTCCGACCTGGGCTACTTCGGTAGGCTAAGGGAGAAGAGATAACAGATACTTCGGTATCTG<br>TTATCTGTTTTTTTTCAACAGATAGCCGCGTTCGCGCGGCTATCTGTTTTTTTTAGGCTAGGTGGAGGCTCAGTGATGAGATCGATAACGCGCCGGAAGAAAAAGCT<br>CGTGGTATCACCATCAAATTCTAATACGACTCACTATAGGAGCGGTAGTTCAGTCGGTTAGAATACCTGCCTGTCACGCAGGGGGTCGCGGGTTCGAGTCCCGTCC<br>GTTCCGCCACGCGGGGTGGAGCAGCCTGGTAGCTCGTCGGGCTCATAACCCGAAGATCGTCGGTTCAAATCCGGCCCCCGCAACCAGGTGGCTATAGCTCAGTTGG<br>TAGAGCCCTGGATTGTGATTCCAGTTGTCGTGGGTTCGAATCCCATTAGCCACCCCAGGGTCGTTAGCTCAGTTGGTAGAGCAGTTGACTCTTAATCAATTGGTCGC<br>AGGTTCGAATCCTGCACGACCCACCAGCGTCCGTAGCTCAGTTGGTTAGAGCACCACCTTGACATGGTGGGGGTCGGTGGTTCGAGTCCACTCGGACGCACCAGGG<br>TCGGCATGGCATCTCCACCTCCTCGCGGTCCGACCTGGGCTACTTCGGTAGGCTAAGGGAGAAGAGATAACAGATACTTCGGTATCTGTTATCTGTTTTTTTTCAAC<br>AGATAGCCGCGTTCGCGCGGCTATCTGTTTTTTTTAGGCTAGGTGGAGGCTCAGTGATGACACTTCTCACGTTGAATACGACACCCCGACCCGTCACTACGCACAAT<br>TCTAATACGACTCACTATAGGTGAGGTGTCCGAGTGGCTGAAGGAGCACGCCTGGAAAGTGTGTATACGGCAACGTATCGGGGGTTCGAATCCCCCCCTCACCGCC<br>AGGCTACGTAGCTCAGTTGGTTAGAGCACATCACTCATAATGATGGGGTCACAGGTTCGAATCCCGTCGTAGCCACCAGCCCGGATAGCTCAGTCGGTAGAGCAGG<br>GGATTGAAAATCCCCGTGTCCTTGGTTCGATTCCGAGTCCGGGCACCAGCTGATATGGCTCAGTTGGTAGAGCGCACCCTTGGTAAGGGTGAGGTCCCCAGTTCGA<br>CTCTGGGTATCAGCACCAGGGTCGGCATGGCATCTCCACCTCCTCGCGGTCCGACCTGGGCTACTTCGGTAGGCTAAGGGAGAAGAGATAACAGATACTTCGGTAT<br>CTGTTATCTGTTTTTTTTCAACAGATAGCCGCGTTCGCGCGGCTATCTGTTTTTTTTAGGCTAGGTGGAGGCTCAGTGATGACGTAGACTGCCCGGGGCACGCCGAC<br>TATGTTAAAAACATGATCAAATTCTAATACGACTCACTATAGGGATAATACAACGGTTTCCCATATATCGGCACGTAGCGCAGCCTGGAAGCGCACCGTCATGGGG<br>TGACGGGGGTCGGAGGTTCAAATCCTCTCGTGCCGACCAGGGCTTGTAGCTCAGATGGTTAGAGCGCACCCCTGATAAGGGTGAGGTCGGTGGTTCAAGTCCACTC<br>AGGCCTACCAGTCCCCTTCGTCTAGAGGCCCAGGACACCGCCCTCTCACGGCGGTAACAGGGGTTCGAATCCCCTAGGGGACGCCAGCCTCTGTAGTTCAGTGGGT<br>AGAACGGCGGACTGTTAATCCGTATGTCACTGGTTCGAGTCCAGTCAGAGGCGCCAGGGTCGGCATGGCATCTCCACCTCCTCGCGGTCCGACCTGGGCTACTTCG<br>GTAGGCTAAGGGAGAAGAGATAACAGATACTTCGGTATCTGTTATCTGTTTTTTTTCAACAGATAGCCGCGTTCGCGCGGCTATCTGTTTTTTTTGCTGAAAGGAGG<br>AACTATATCCGGATAACCTGGCAGGCTAGGTGGAGGCTCAGTGATGATAAGTCTGCGATGGTGGATGCATGTGTCATGGTCATAGCTGTTTCCTGTGTGAAATTGTT<br>ATCCGCTCAGAGGGCACAATCCTATTCCGCGCTATCCGACAATCTCCAAGACATTAGGTGGAGTTCAGTTCGGCGTATGGCATATGTCGCTGGAAAGAACATGTGA<br>GCAAAAGGCCAGCAAAAGGCCAGGAACCGTAAAAAGGCCGCGTTGCTGGCGTTTTTCCATAGGCTCCGCCCCCCTGACGAGCATCACAAAAATCGACGCTCAAGT<br>CAGAGGTGGCGAAACCCGACAGGACTATAAAGATACCAGGCGTTTCCCCCTGGAAGCTCCCTCGTGCGCTCTCCTGTTCCGACCCTGCCGCTTACCGGATACCTGT<br>CCGCCTTTCTCCCTTCGGGAAGCGTGGCGCTTTCTCATAGCTCACGCTGTAGGTATCTCAGTTCGGTGTAGGTCGTTCGCTCCAAGCTGGGCTGTGTGCACGAACCC<br>CCCGTTCAGCCCGACCGCTGCGCCTTATCCGGTAACTATCGTCTTGAGTCCAACCCGGTAAGACACGACTTATCGCCACTGGCAGCAGCCACTGGTAACAGGATTA<br>GCAGAGCGAGGTATGTAGGCGGTGCTACAGAGTTCTTGAAGTGGTGGCCTAACTACGGCTACACTAGAAGAACAGTATTTGGTATCTGCGCTCTGCTGAAGCCAGT<br>TACCTTCGGAAAAAGAGTTGGTAGCTCTTGATCCGGCAAACAAACCACCGCTGGTAGCGGTGGTTTTTTTGTTTGCAAGCAGCAGATTACGCGCAGAAAAAAAGGA<br>TCTCAAGAAGATCCTTTGATCTTTTCTACGGGGTCTGACGCTCTATTCAACAAAGCCGCCGTCCCGTCAAGTCAGCGTAAATGGGTAGGGGGCTTCAAATCGTCCTC<br>GTGATACCAATTCGGAGCCTGCTTTTTTGTACAAACTTGTTGATAATGGCAATTCAAGGATCTTCACCTAGATCCTTTTAAATTAAAAATGAAGTTTTAAATCAATCT<br>AAAGTATATATGAGTAAACTTGGTCTGACAGTTACCAATGCTTAATCAGTGAGGCACCTATCTCAGCGATCTGTCTATTTCGTTCATCCATAGTTGCCTGACTCCCC<br>GTCGTGTAGATAACTACGATACGGGAGGGCTTACCATCTGGCCCCAGTGCTGCAATGATACCGCGAGAGCCACGCTCACCGGCTCCAGATTTATCAGCAATAAACC<br>AGCCAGCCGGAAGGGCCGAGCGCAGAAGTGGTCCTGCAACTTTATCCGCCTCCATCCAGTCTATTAATTGTTGCCGGGAAGCTAGAGTAAGTAGTTCGCCAGTTAA<br>TAGTTTGCGCAACGTTGTTGCCATTGCTACAGGCATCGTGGTGTCACGCTCGTCGTTTGGTATGGCTTCATTCAGCTCCGGTTCCCAACGATCAAGGCGAGTTACAT<br>GATCCCCCATGTTGTGCAAAAAAGCGGTTAGCTCCTTCGGTCCTCCGATCGTTGTCAGAAGTAAGTTGGCCGCAGTGTTATCACTCATGGTTATGGCAGCACTGCAT<br>AATTCTCTTACTGTCATGCCATCCGTAAGATGCTTTTCTGTGACTGGTGAGTACTCAACCAAGTCATTCTGAGAATAGTGTATGCGGCGACCGAGTTGCTCTTGCCC<br>GGCGTCAATACGGGATAATACCGCGCCACATAGCAGAACTTTAAAAGTGCTCATCATTGGAAAACGTTCTTCGGGGCGAAAACTCTCAAGGATCTTACCGCTGTTG<br>AGATCCAGTTCGATGTAACCCACTCGTGCACCCAACTGATCTTCAGCATCTTTTACTTTCACCAGCGTTTCTGGGTGAGCAAAAACAGGAAGGCAAAATGCCGCAA<br>AAAAGGGAATAAGGGCGACACGGAAATGTTGAATACTCATACTCTTCCTTTTTCAATATTATTGAAGCATTTATCAGGGTTATTGTCTCATGAGCGGATACATATTT<br>GAATGTATTTAGAAAAATAAACAAATAGGGGTTCCGCGCACATTTCCCCGAAAAGTGCCAGATACCTGAAACAAAACCCATCGTACGGCCAAGGAAGTCTCCAAT<br>AACTGTGATCCACCACAAGCGCCAGGGTTTTCCCAGTCACGACGTTGTAAAACGACGGCCAGTCATGCATAATCCGCACGCATCTGGAATAAGGAAGTGCCATTCC<br>GCCTGACCT |
| pUC_M1<br>*M1RNA | AGTTCGGTGTAGGTCGTTCGCTCCAAGCTGGGCTGTGTGCACGAACCCCCCGTTCAGCCCGACCGCTGCGCCTTATCCGGTAACTATCGTCTTGAGTCCAACCCGGT<br>AAGACACGACTTATCGCCACTGGCAGCAGCCACTGGTAACAGGATTAGCAGAGCGAGGTATGTAGGCGGTGCTACAGAGTTCTTGAAGTGGTGGCCTAACTACGG<br>CTACACTAGAAGAACAGTATTTGGTATCTGCGCTCTGCTGAAGCCAGTTACCTTCGGAAAAAGAGTTGGTAGCTCTTGATCCGGCAAACAAACCACCGCTGGTAGC<br>GGTGGTTTTTTTGTTTGCAAGCAGCAGATTACGCGCAGAAAAAAAGGATCTCAAGAAGATCCTTTGATCTTTTCTACGGGGTCTGACGCTCAGTGGAACGAAAACT<br>CACGTTAAGGGATTTTGGTCATGAGATTATCAAAAAGGATCTTCACCTAGATCCTTTTAAATTAAAAATGAAGTTTTAAATCAATCTAAAGTATATATGAGTAAACT<br>TGGTCTGACAGTTACCAATGCTTAATCAGTGAGGCACCTATCTCAGCGATCTGTCTATTTCGTTCATCCATAGTTGCCTGACTCCCCGTCGTGTAGATAACTACGATA<br>CGGGAGGGCTTACCATCTGGCCCCAGTGCTGCAATGATACCGCGAGACCCACGCTCACCGGCTCCAGATTTATCAGCAATAAACCAGCCAGCCGGAAGGGCCGAG |

| | |
|---|---|
| | CGCAGAAGTGGTCCTGCAACTTTATCCGCCTCCATCCAGTCTATTAATTGTTGCCGGGAAGCTAGAGTAAGTAGTTCGCCAGTTAATAGTTTGCGCAACGTTGTTGC<br>CATTGCTACAGGCATCGTGGTGTCACGCTCGTCGTTTGGTATGGCTTCATTCAGCTCCGGTTCCCAACGATCAAGGCGAGTTACATGATCCCCCATGTTGTGCAAAA<br>AAGCGGTTAGCTCCTTCGGTCCTCCGATCGTTGTCAGAAGTAAGTTGGCCGCAGTGTTATCACTCATGGTTATGGCAGCACTGCATAATTCTCTTACTGTCATGCCA<br>TCCGTAAGATGCTTTTCTGTGACTGGTGAGTACTCAACCAAGTCATTCTGAGAATAGTGTATGCGGCGACCGAGTTGCTCTTGCCCGGCGTCAATACGGGATAATAC<br>CGCGCCACATAGCAGAACTTTAAAAGTGCTCATCATTGGAAAACGTTCTTCGGGGCGAAAACTCTCAAGGATCTTACCGCTGTTGAGATCCAGTTCGATGTAACCC<br>ACTCGTGCACCCAACTGATCTTCAGCATCTTTTACTTTCACCAGCGTTTCTGGGTGAGCAAAAACAGGAAGGCAAAATGCCGCAAAAAAGGGAATAAGGGCGACA<br>CGGAAATGTTGAATACTCATACTCTTCCTTTTTCAATATTATTGAAGCATTTATCAGGGTTATTGTCTCATGAGCGGATACATATTTGAATGTATTTAGAAAAATAAA<br>CAAATAGGGGTTCCGCGCACATTTCCCCGAAAAGTGCCACCTGACGTCTAAGAAACCATTATTATCATGACATTAACCTATAAAAATAGGCGTATCACGAGGCCCT<br>TTCGTCTCGCGCGTTTCGGTGATGACGGTGAAAACCTCTGACACATGCAGCTCCCGGAGACGGTCACAGCTTGTCTGTAAGCGGATGCCGGGAGCAGACAAGCCCG<br>TCAGGGCGCGTCAGCGGGTGTTGGCGGGTGTCGGGGCTGGCTTAACTATGCGGCATCAGAGCAGATTGTACTGAGAGTGCACCATATGCGGTGTGAAATACCGCA<br>CAGATGCGTAAGGAGAAAATACCGCATCAGGCGCCATTCGCCATTCAGGCTGCGCAACTGTTGGGAAGGGCGATCGGTGCGGGCCTCTTCGCTATTACGCCAGCTG<br>GCGAAAGGGGGATGTGCTGCAAGGCGATTAAGTTGGGTAACGCCAGGGTTTTCCCAGTCACGACGTTGTAAAACGACGGCCAGTGAATTCTAATACGACTCACTAT<br>AGAAGCTGACCAGACAGTCGCCGCTTCGTCGTCGTCCTCTTCGGGGGAGACGGGCGGAGGGGAGGAAAGTCCGGGCTCCATAGGGCAGGGTGCCAGGTAACGCCT<br>GGGGGGGAAACCCACGACCAGTGCAACAGAGAGCAAACCGCCGATGGCCCGCGCAAGCGGGATCAGGTAAGGGTGAAAGGGTGCGGTAAGAGCGCACCGCGCG<br>GCTGGTAACAGTCCGTGGCACGGTAAACTCCACCCGGAGCAAGGCCAAATAGGGGTTCATAAGGTACGGCCCGTACTGAACCCGGGTAGGCTGCTTGAGCCAGTG<br>AGCGATTGCTGGCCTAGATGAATGACTGTCCACGACAGAACCCGGCTTATCGGTCAGTTTCACCTGATTTACGTCATCCGGATCCTCTAGAGTCGACCTGCAGGCAT<br>GCAAGCTTGGCGTAATCATGGTCATAGCTGTTTCCTGTGTGAAATTGTTATCCGCTCACAATTCCACACAACATACGAGCCGGAAGCATAAAGTGTAAAGCCTGGG<br>GTGCCTAATGAGTGAGCTAACTCACATTAATTGCGTTGCGCTCACTGCCCGCTTTCCAGTCGGGAAACCTGTCGTGCCAGCTGCATTAATGAATCGGCCAACGCGCG<br>GGGAGAGGCGGTTTGCGTATTGGGCGCTCTTCCGCTTCCTCGCTCACTGACTCGCTGCGCTCGGTCGTTCGGCTGCGGCGAGCGGTATCAGCTCACTCAAAGGCGGT<br>AATACGGTTATCCACAGAATCAGGGGATAACGCAGGAAAGAACATGTGAGCAAAAGGCCAGCAAAAGGCCAGGAACCGTAAAAAGGCCGCGTTGCTGGCGTTTT<br>TCCATAGGCTCCGCCCCCCTGACGAGCATCACAAAAATCGACGCTCAAGTCAGAGGTGGCGAAACCCGACAGGACTATAAAGATACCAGGCGTTTCCCCCTGGAA<br>GCTCCCTCGTGCGCTCTCCTGTTCCGACCCTGCCGCTTACCGGATACCTGTCCGCCTTTCTCCCTTCGGGAAGCGTGGCGCTTTCTCATAGCTCACGCTGTAGGTATC<br>TC |

**Table S4. Sequences of the 21 tRNAs used in this study**

| tRNA | Sequence |
|---|---|
| tRNA$^{Ala}$GGC | GGGGCUAUAGCUCAGCUGGGAGAGCGCUUGCAUGGCAUGCAAGAGGUCAGCGGU UCGAUCCCGCUUAGCUCCACCA |
| tRNA$^{Arg}$CCG | GCGCCCGUAGCUCAGCUGGAUAGAGCGCUGCCCUCCGGAGGCAGAGGUCUCAGGU UCGAAUCCUGUCGGGCGCGCCA |
| tRNA$^{Asp}$GUC | GGAGCGGUAGUUCAGUCGGUUAGAAUACCUGCCUGUCACGCAGGGGGUCGCGGG UUCGAGUCCCGUCCGUUCCGCCA |
| tRNA$^{Cys}$GCA | GGCGCGUUAACAAAGCGGUUAUGUAGCGGAUUGCAAAUCCGUCUAGUCCGGUUC GACUCCGGAACGCGCCUCCA |
| tRNA$^{Gly}$GCG | GCGGGAAUAGCUCAGUUGGUAGAGCACGACCUUGCCAAGGUCGGGGUCGCGAGU UCGAGUCUCGUUUCCCGCUCCA |
| tRNA$^{Glu}$CUC | GUCCCCUUCGUCUAGAGGCCCAGGACACCGCCCUCUCACGGCGGUAACAGGGGUU CGAAUCCCCUAGGGGACGCCA |
| tRNA$^{His}$GUG | GGUGGCUAUAGCUCAGUUGGUAGAGCCCUGGAUUGUGAUUCCAGUUGUCGUGGG UUCGAAUCCCAUUAGCCACCCCA |
| tRNA$^{Leu}$CAG | GCGAAGGUGGCGGAAUUGGUAGACGCGCUAGCUUCAGGUGUUAGUGUCCUUACG GACGUGGGGGUUCAAGUCCCCCCCCUCGCACCA |
| tRNA$^{Lys}$CUU | GGGUCGUUAGCUCAGUUGGUAGAGCAGUUGACUCUUAAUCAAUUGGUCGCAGGU UCGAAUCCUGCACGACCCACCA |
| tRNA$^{mMet}$CAU | GGCUACGUAGCUCAGUUGGUUAGAGCACAUCACUCAUAAUGAUGGGGUCACAGG UUCGAAUCCCGUCGUAGCCACCA |
| tRNA$^{Phe}$GAA | GCCCGGAUAGCUCAGUCGGUAGAGCAGGGGAUUGAAAAUCCCCGUGUCCUUGGU UCGAUUCCGAGUCCGGGCACCA |
| tRNA$^{Ser}$GGA | GGUGAGGUGUCCGAGUGGCUGAAGGAGCACGCCUGGAAAGUGUGUAUACGGCAA CGUAUCGGGGGUUCGAAUCCCCCCCUCACCGCCA |
| tRNA$^{Thr}$GGU | GCUGAUAUGGCUCAGUUGGUAGAGCGCACCCUUGGUAAGGGUGAGGUCCCCAGU UCGACUCUGGGUAUCAGCACCA |
| tRNA$^{Tyr}$GUA | GGUGGGGUUCCCGAGCGGCCAAAGGGAGCAGACUGUAAAUCUGCCGUCACAGAC UUCGAAGGUUCGAAUCCUUCCCCCACCACCA |
| tRNA$^{Val}$GAC | GCGUCCGUAGCUCAGUUGGUUAGAGCACCACCUUGACAUGGUGGGGGUCGGUGG UUCGAGUCCACUCGGACGCACCA |
| tRNA$^{Asn}$GUU | UCCUCUGUAGUUCAGUCGGUAGAACGGCGGACUGUUAAUCCGUAUGUCACUGGU UCGAGUCCAGUCAGAGGAGCCA |
| tRNA$^{Gln}$CUG | UGGGGUAUCGCCAAGCGGUAAGGCACCGGAUUCUGAUUCCGGCAUUCCGAGGUU CGAAUCCUCGUACCCCAGCCA |
| tRNA$^{Ile}$GAU | AGGCUUGUAGCUCAGGUGGUUAGAGCGCACCCCUGAUAAGGGUGAGGUCGGUGG UUCAAGUCCACUCAGGCCUACCA |
| tRNA$^{fMet}$CAU | CGCGGGGUGGAGCAGCCUGGUAGCUCGUCGGGCUCAUAACCCGAAGAUCGUCGG UUCAAAUCCGGCCCCCGCAACCA |
| tRNA$^{Pro}$GGG | CGGCACGUAGCGCAGCCUGGUAGCGCACCGUCAUGGGGUGUCGGGGGUCGGAGG UUCAAAUCCUCUCGUGCCGACCA |
| tRNA$^{Trp}$CCA | AGGGGCGUAGUUCAAUUGGUAGAGCACCGGUCUCCAAAACCGGGUGUUGGGAGU UCGAGUCUCUCCGCCCCUGCCA |
| tRNA$^{Ile}$GAU_v2 | AGGCUUGUAGCUCAGAUGGUUAGAGCGCACCCCUGAUAAGGGUGAGGUCGGUGG UUCAAGUCCACUCAGGCCUACCA |
| tRNA$^{Asn}$GUU_v2 | UCCUCUGUAGUUCAGUGGGUAGAACGGCGGACUGUUAAUCCGUAUGUCACUGGU UCGAGUCCAGUCAGAGGAGCCA |
| tRNA$^{Pro}$GGG_v2 | CGGCACGUAGCGCAGCCUGGAAGCGCACCGUCAUGGGGUGACGGGGGUCGGAGG UUCAAAUCCUCUCGUGCCGACCA |

**Table S5. Standard composition of the tRNA-free PURE system (tfPURE)**

| Component | Concentration | Component | Concentration |
|---|---|---|---|
| Initiation Factor 1 | 25 µM | Tryptophanyl-tRNA Synthetase | 28 nM |
| Initiation Factor 2 | 1.0 µM | Tyrosyl-tRNA Synthetase | 0.15 µM |
| Initiation Factor 3 | 4.9 µM | Valyl-tRNA Synthetase | 17 nM |
| Elongation Factor G | 1.1 µM | Methionyl-tRNA Formyltransferase | 0.59 µM |
| Elongation Factor Tu | 80 µM | Myokinase | 1.4 µM |
| Elongation Factor Ts | 3.3 µM | Creatine kinase | 0.25 µM |
| Release Factor 1 | 49 nM | Nucleoside diphosphate kinase | 16 nM |
| Release Factor 2 | 48 nM | Pyrophosphatase | 41 nM |
| Release Factor 3 | 0.17 µM | Trigger Factor | 1.0 µM |
| Ribosome Recycling Factor | 3.9 µM | E. coli DEAH type RNA helicase A | 63 nM |
| Alanyl-tRNA Synthetase | 0.73 µM | Ribosome | 1.0 µM |
| Arginyl-tRNA Synthetase | 31 nM | Tyrosine | 0.30 mM |
| Asparaginyl-tRNA Synthetase | 0.42 µM | Cysteine | 0.30 mM |
| Asparagyl-tRNA Synthetase | 0.12 µM | 18 other amino acids | 0.36 mM |
| Cysteinyl-tRNA Synthetase | 24 nM | ATP | 0.38* mM |
| Glutaminyl-tRNA Synthetase | 60 nM | GTP | 0.25* mM |
| Glutamyl-tRNA Synthetase | 0.23 µM | CTP | 0.13* mM |
| Glycyl-tRNA Synthetase | 86 nM | UTP | 0.13* mM |
| Histidyl-tRNA Synthetase | 85 nM | N-2-hydroxyethylpiperazine-N'-2-ethanesulfonic acid (pH7.6) | 0.10 M |
| Isoleucyl-tRNA Synthetase | 0.37 µM | Glutamic acid potassium salt | 70 mM |
| Leucyl-tRNA Synthetase | 41 nM | Spermidine | 0.375* mM |
| Lysyl-tRNA Synthetase | 0.12 µM | Creatine phosphate | 25 mM |
| Methionyl-tRNA Synthetase | 0.11 µM | Dithiothreitol | 6 mM |
| Phenylalanyl-tRNA Synthetase | 0.13 µM | 10-formyl-5,6,7,8-tetrahydro folic acid | 10 µg/mL |
| Prolyl-tRNA Synthetase | 0.17 µM | Yeast inorganic pyrophosphatase (NEB) | 0.2 units/mL |
| Seryl-tRNA Synthetase | 78 nM | RNase Plus Inhibitor (Promega) | 0.1 U/µL |
| Threonyl-tRNA Synthetase | 84 nM | T7 RNAP (Takara) | 0.42* U/µL |

*Concentrations in the composition A are shown. Concentrations in other compositions are shown in Table S6.

**Table S6. Reaction compositions of tfPURE systems under different experimental conditions**

| Component | PURE system composition | | |
|---|---|---|---|
| | A | B | C |
| NTP (mM) | 0.88 | 3.1 | 7.4 |
| $Mg(OAc)_2$ (mM) | 7.9 | 16 | 16 |
| Spermidine (mM) | 0.38 | 0.75 | 0.75 |
| T7 RNAP (U/μL) | 0.42 | 1.7 | 3.4 |
